# Markov State Models of Proton- and Gate-Dependent Activation in a Pentameric Ligand-Gated Ion Channel

**DOI:** 10.1101/2021.03.12.435097

**Authors:** Cathrine Bergh, Stephanie A. Heusser, Rebecca J. Howard, Erik Lindahl

## Abstract

Ligand-gated ion channels conduct currents in response to chemical stimuli, mediating electrochemical signaling in neurons and other excitable cells. For many channels the mechanistic details of gating remain unclear, partly due to limited structural data and simulation timescales. Here, we used enhanced sampling to simulate the pH-gated channel GLIC, and construct Markov state models (MSMs) of gating transitions. Consistent with new functional recordings reported here in oocytes, our analysis revealed differential effects of protonation and mutation on free-energy wells. Clustering of closed-versus open-like states enabled estimation of open probabilities and transition rates in each condition, while higher-order clustering affirmed conformational trends in gating. Furthermore, our models uncovered state- and protonation-dependent symmetrization among subunits. This demonstrates the applicability of MSMs to map energetic and conformational transitions between ion-channel functional states, and how they correctly reproduce shifts upon activation or mutation, with implications for modeling neuronal function and developing state-selective drugs.

## Introduction

The family of pentameric ligand-gated ion channels (pLGICs), also known as Cys-loop receptors, controls electrochemical signal transduction in numerous tissues and cell types, from bacteria to humans. A rapid cycling between conducting and nonconducting conformations in response to chemical stimuli, such as neurotransmitter binding or changes in pH, is fundamental to their function. These channels are often found in the postsynaptic membrane of neurons and undergo allosteric conformational changes, where the pore in the integral transmembrane domain (TMD) opens for ion conduction upon selective neurotransmitter binding in the extracellular domain (ECD). Prokaryotic homologs, such as the proton-gated channel GLIC from the cyanobacterium *Gloeobacter violaceus*, share many topological features with eukaryotic pLGICs, and have been proposed to follow similar gating pathways (Figure 1A-D). For any pLGIC, channel function can ultimately be explained by its free energy landscape, where different state populations shift upon activation through agonist binding. Understanding these landscapes and how the free energies either of stable states or barriers change upon ligand-binding or mutation is thus crucial for full understanding of the gating process, with applications including the development of state-dependent drugs for better treatment of diseases related to channel malfunction.

**Figure 1:**
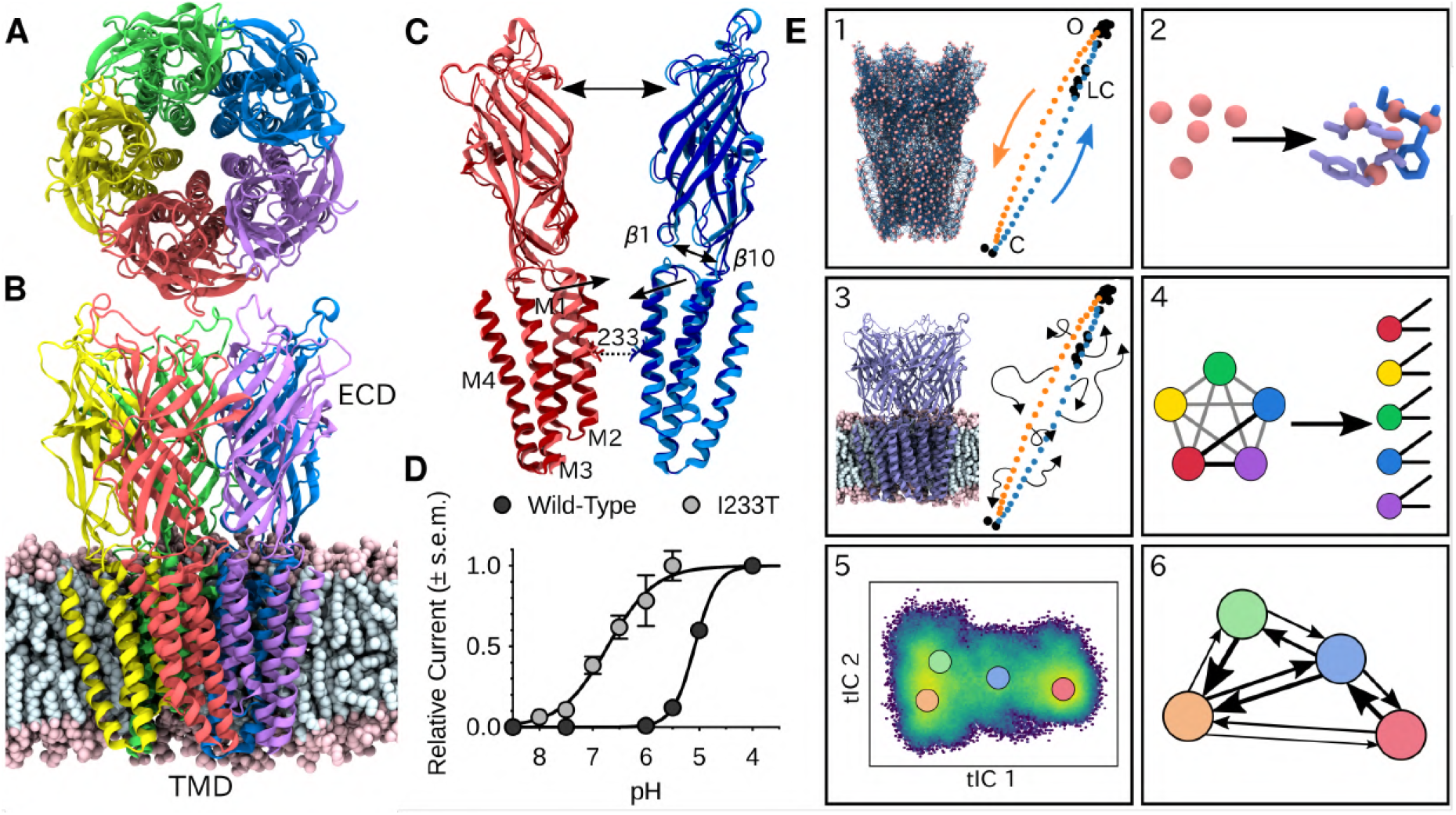
Global architecture of GLIC, electrophysiology data and computational methodology. GLIC in an open conformation shown from (A) the top and (B) the side in a POPC lipid bilayer. (C) Two opposing subunits highlighting the pore of the channel. Light colors represent the open conformation (PDB ID 4HFI) and dark colors the closed one (PDB ID 4NPQ). Arrows indicate important gating motions - the tilting of the M2 helices, beta expansion and ECD spread. Residue I233 at the 9’ position was mutated in both simulations and electrophysiology experiments. (D) Electrophysiology data for wild-type GLIC and the gain-of-function I233T mutation. (E) Simulation methodology: the eBDIMS method provides (1) coarse-grained seed structures along the transition pathway followed by (2) reconstruction of the atomistic detail. Atomistic structures were then (3) embedded in lipid bilayers and massively parallel unrestrained MD launched. Analysis involves (4) a feature transformation to account for the symmetry of the pentamer, followed by (5) dimensionality reduction with tICA and (6) MSM construction.

Recent advances in structural biology have enabled a steady increase in the number of available pLGIC structures. The GLIC model system is notable in this regard, accounting for over 40% of pLGIC entries in the protein data bank, including apparent closed and open states. However, the conformational diversity of these states is highly limited, leading to crude representations of the free-energy landscapes from experimental structures alone. Computational methods like molecular dynamics (MD) simulations can be used to sample more of the conformational landscape, and several studies have been conducted on GLIC to study short-timescale motions [1–12]. Still, due to the large system size and relatively long timescales of the gating transitions, in practice it has not been feasible to sample complete gating transitions, especially if ligandbinding and unbinding events are involved [13-19]. To bridge this gap, various enhanced sampling methods can be used, often by the application of biasing forces or presumed reaction coordinates. For instance, application of the string method with swarms of trajectories recently enabled the identification of local rearrangements in channel closure, including contraction of the upper pore, loosening of β-strand contacts in the lower ECD, and general expansion of the upper ECD [20]. This provides precious information of structural rearrangements, but the choice of collective variables in combination with the short ps-timescale of individual simulations may influence what motions are sampled.

Markov state models (MSMs) show promise in modeling the thermodynamics and kinetics of biological systems without making prior mechanistic assumptions [21–23]. By counting transitions, multiple simulations can effectively be stitched together to capture processes on timescales longer than any individual simulation, and probing more biologically relevant dynamics. However, in principle MSMs first require the whole equilibrium distribution to be sampled, which is difficult to achieve by starting simulations from experimental structures alone, since the stability required for crystal packing or cryo-EM data processing results in structures mostly representing metastable states. Simulations started from these states then usually remain confined to the energy well for long time periods. In contrast, seeding simulations at regions in the free energy landscape that are not necessarily metastable enhance sampling without introducing any biasing forces, and the actual sampling is performed without limiting the system to any particular reaction coordinate.

Here, we have used such enhanced seeding approaches combined with MSMs to sample the GLIC opening-closing transition. Both the wild-type and a variant with a mutation of the main hydrophobic gate, which we also showed to yield gain-of-function similar to human homologs [24, 25], were simulated in resting or activating conditions (neutral or low pH). From the resulting trajectories we were able to construct MSMs of the complete ion channel, quantitatively map free energy landscapes, extract molecular details regarding the gating mechanisms, and show how the gating shift observed electrophysiology recordings of the wild-type and variant is modeled correctly by the MSMs. Additionally, we present new evidence for the role of symmetry in gating.

## Results

### Enhanced Sampling Enables MSM Construction of GLIC Gating

To shed light on the gating mechanisms, we combined enhanced sampling of MD simulations at resting and activating pH with Markov state modeling. X-ray structures of GLIC crystallized at pH 7 (PDB ID 4NPQ) and pH 4 (PDB ID 4HFI) have been reported to represent closed and open states, respectively [26, 27] (Figure 1C). Because unbiased molecular dynamics simulations were not expected to thoroughly sample the activation process, we seeded simulations along the presumed gating transition which enabled simulations to run in a massively parallel fashion. To achieve this, the starting structures were simplified to C*α* traces, and used to drive elastic network-driven Brownian dynamics (eBDIMS) [28, 29] to generate two coarse-grained pathways: one from the closed to the open X-ray structure, and one in the opposite direction (Figure 1E1). Following side-chain reconstruction (Figure 1E2), this approach resulted in a set of 50 initial models interspersed in principal component space between the open and closed X-ray structure clusters, all with standard titration states representing neutral pH (deprotonated) that in experiments should result in a closed channel. In a duplicate set of initial models, a subset of acidic residues was modified to the most probable titration state under activating conditions (protonated), as previously described [30]. Each resulting model was then subjected to unrestrained simulation in a palmitoyloleoylphosphatidylcholine (POPC) lipid bilayer for over 1 *μ*s, producing 120 *μ*s total sampling in the two conditions (Figure 1E3). Among simulations performed at each pH, we performed a feature transformation to account for the symmetry of the homopentameric protein (Figure 1E4), followed by dimensionality reduction by time-lagged independent component analysis (tICA) (Figure 1E5). Further clustering in the resulting tICA space yielded kinetically meaningful states that could be used for MSM construction (Figure 1E6), validated by convergence of the main transition timescale (Figure S4).

To assess the ability of this computational approach to predict functional properties, we introduced the gain-of-function mutation I233T, located at the midpoint of the GLIC transmembrane pore (Figure 1C).

This position, 9’ counting from the intracellular side, has been shown to constitute a hydrophobic gate that critically influences conduction properties in GLIC as in other family members [31]. Indeed, we confirmed by two-electrode voltage-clamp electrophysiology recordings in *Xenopus oocytes* that the polar threonine substitution enhanced proton sensitivity, shifting half-maximal activation by more than one pH unit, and producing moderately conductive channels even at neutral pH (Figure 1D). This I233T substitution was introduced into additional replicate sets of initial models in both deprotonated and protonated states as described above. The resulting mutant was prepared, simulated, transformed, and analyzed in the same way as the wild-type channels, achieving convergence on similar timescales (Figure S4). Thus we were able to produce four independent, statistically validated, MSMs representing four distinct combinations of pH and hydrophobic-gate variations.

### Free-Energy Landscapes Capture Effects of Protonation and Mutation

Free energy landscapes obtained from the first eigenvector of each MSM clearly distinguished closed- and open-like regions in all four conditions (Figure 2). Along the first tIC - presumed to represent the slowest transition - the closed X-ray structure consistently projected to the lowest free energy minimum in the landscape, suggesting a predominant population of closed channels under both deprotonated and protonated conditions. Due, in part, to its low conductance in single-channel recordings [32], the open probability of GLIC is not well established; however, recent cryo-electron microscopy studies support a predominance of nonconducting states even at pH below 4 [33]. In deprotonated conditions, the I233T substitution created a well-defined second local minimum in free energy along tIC1, centered near the open X-ray structures (Figure 2A, 2C). Protonated conditions further deepened this secondary minimum for both wild-type and I233T variants (Figure 2B, 2D). Despite topological differences, the distribution of closed versus open X-ray structures was comparable along tIC1 in all four conditions, suggesting a common component for this slowest transition. Several so-called locally closed structures (e.g. PDB ID 3TLS) - featuring a closed-like TMD but open-like ECD - projected to a region intermediate along tIC1, but within the broad closed-state free energy well, suggesting this component primarily distinguishes TMD rather than ECD state. Conversely, a lipid-bound state (PDB ID 5J0Z) - in which the pore is expanded at the outward vestibule, though nonconductive at the hydrophobic gate - clustered with open channels. In contrast to tIC1, tIC2 did not distinguish X-ray structures in a consistent manner; higher values of this component may represent conformations sampled primarily in MD simulations.

**Figure 2:**
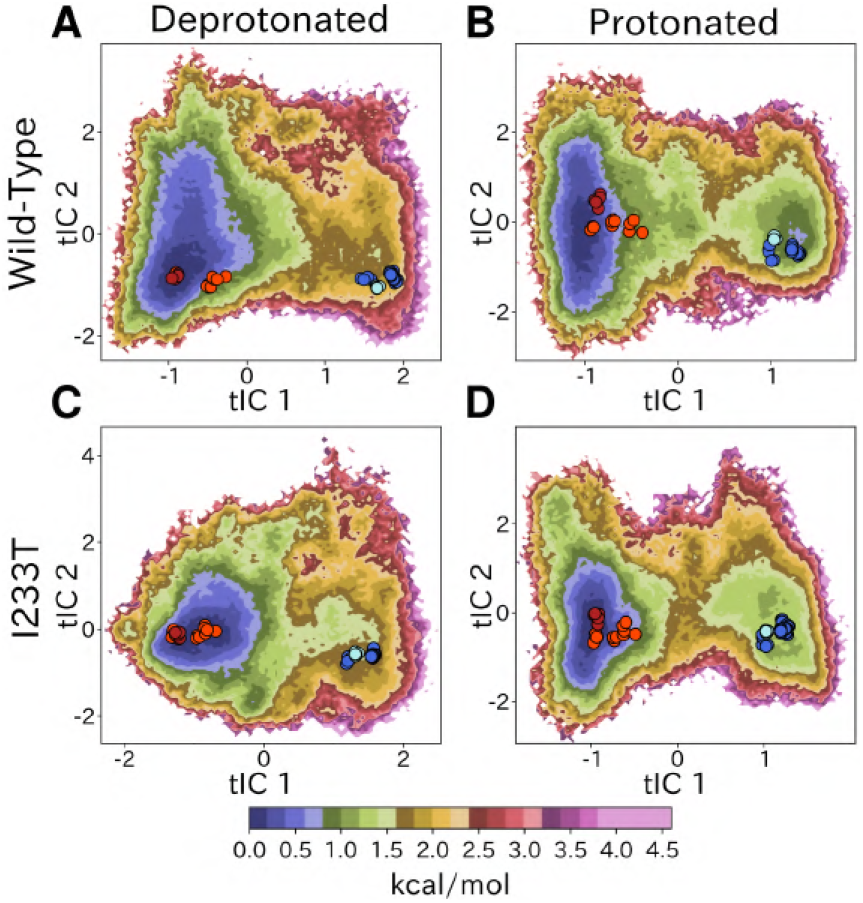
Free energy landscapes capture shifts upon protonation and mutation. Free energy landscapes projected onto the first two tICA coordinates for (A) deprotonated wild-type, (B) protonated wild-type, (C) deprotonated I233T mutant and (D) protonated I233T mutant. Dots indicate experimental structures, with red representing closed states (PDB ID 4NPQ), orange locally closed conformations (PDB IDs 3TLS, 5MUO, 4NPP(B)), light blue modulated states (PDB ID 5J0Z) and blue open states (PDB IDs 4HFI, 3EAM, 3P4W, 4IL4, 3UU5, 4NPP(A)). A sub-maximal amount of the channels are in the open state at protonated conditions, and the closed state is destabilized for the I233T mutant at deprotonated conditions.

### Kinetic Clustering Distinguishes Metastable Open and Closed States

To quantify population shifts in the evident closed- and open-like regions of the tIC landscape, we constructed more coarse-grained MSMs. By clustering according to the first dynamical MSM eigenvector, each free energy landscape could be divided into two macrostates corresponding to closed and open structures, respectively (Figure 3A). In all but the deprotonated wild-type landscape, an evident free energy barrier indicated these states to be metastable (Figure 2). Based on this clustering, and consistent with the qualitative comparisons above, the fractional population of open-like macrostates moderately increased (6% to 12%) upon I233T substitution under deprotonated conditions, and increased further for both variants upon protonation (to 17% and 20% for wild-type and I233T, respectively) (Figure 3B). MSM kinetics indicated the I233T mutation accelerated opening transitions, decreasing the closed–open mean first-passage time by over one third (8.9 to 5.8 *μ*s); conversely, protonation appeared to slow closing transitions, increasing open–closed times nearly 3-fold in both conditions (0.5 to 1.4 *μ*s and 0.6 to 1.7 *μ*s for wild-type and I233T, respectively) (Figure 3C).

**Figure 3:**
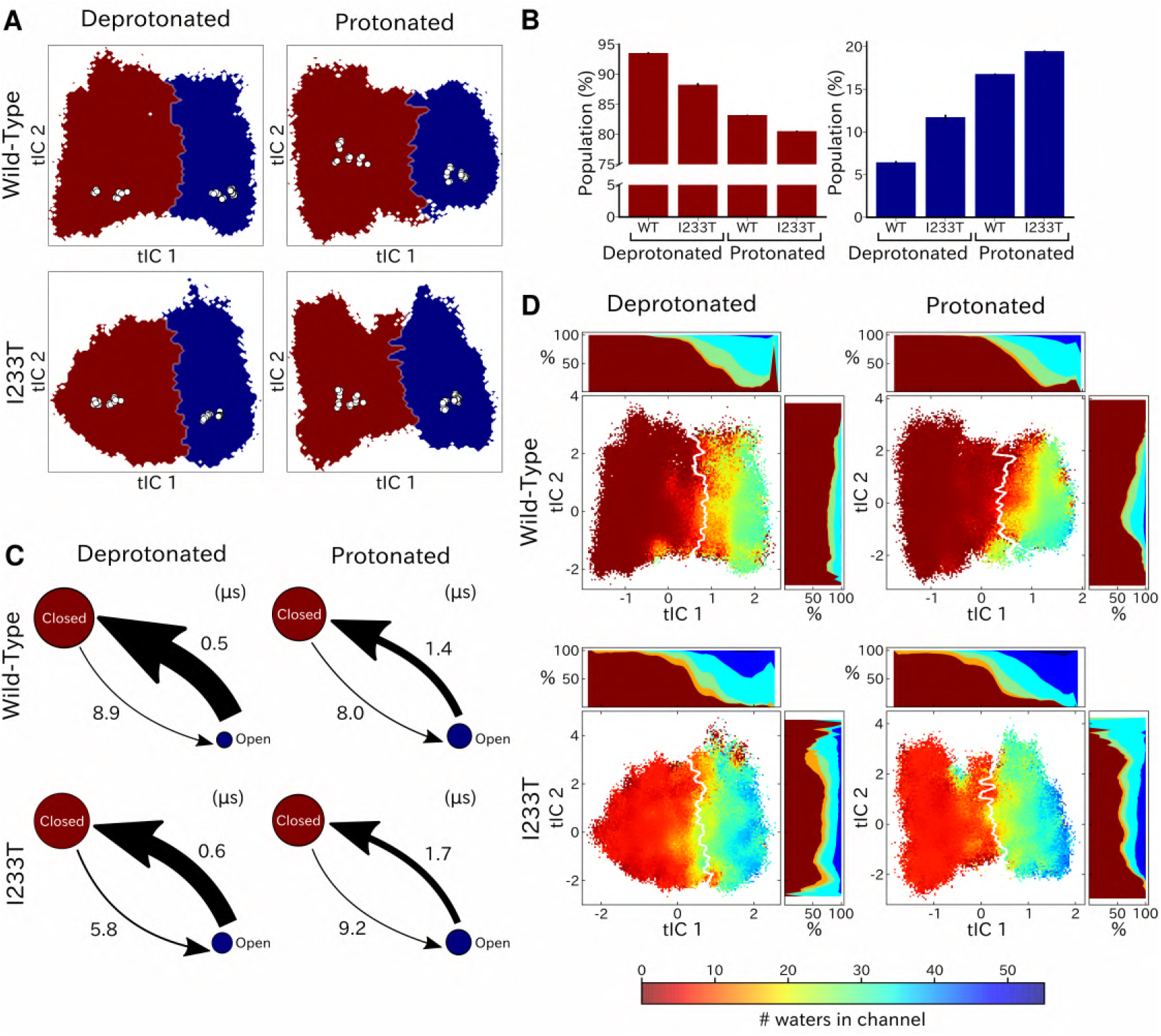
Two-state clustering distinguishes metastable open and closed states. (A) Two metastable states separated by the main free energy barrier, where red represents closed-like states and blue open-like states. (B) Populations of the closed (red) and open (blue) macrostate. Both protonated conditions give the highest values of open probability, *P_O_*, while deprotonated wild-type has the lowest. Deprotonated I233T yields an intermediate level *P_O_*, indicating destabilization of the closed state. For GLIC, absolute values of *P_O_* have not previously been determined, but relative differences agree with our electrophysiology recordings in Figure 1. (C) Transition rates (arrows) with numbers representing mean first passage times. The I233T mutation results in slower transitions between open and closed states at protonated conditions, with the opening transitions significantly faster at deprotonated conditions. (D) Hydration of the transmembrane pore. Side panels show the population of states with different hydration levels. The pore can be seen hydrating or dehydrating when crossing the main free energy barrier (white). The I233T mutation results in higher levels of hydration in both open and closed states.

To further validate the functional annotation of closed- and open-like macrostates, we plotted pore hydration across each free energy landscape (Figure 3D). Under all conditions, the macrostate barrier corresponded to a dramatic shift in hydration levels along tIC1. All regions of tIC space sampled some dehydrated states, likely corresponding to transient, reversible, obstructions observed in individual trajectories (Figures S1–S2). However, protonation increased hydration for both wild-type and I233T variants, particularly at larger (open-like) values of tIC1 (Figure 3D). As expected with a polar residue at the hydrophobic gate, the I233T substitution further increased hydration at all values along tIC1, suggesting more closed-like states might achieve ion conduction in this variant. Thus, qualitative and quantitative comparisons of tIC landscapes supported reproducible state distinctions, and recapitulated functional effects of both protonation and mutation, supporting our model as a reasonable representation of GLIC gating.

### Higher-Order Clustering Reveals Conformational Trends in Gating

To identify conformations along the gating pathway more precisely, we reclustered each dataset according to a larger eigenvector set, obtaining models with five states each (Figure 4A). Note that these states are not necessarily metastable, but this clustering enables further studies of the different regions of the energy landscapes. Conformations corresponding to low values of both tIC1 and tIC2 (state I, Figure 4A) consistently comprised the most populated cluster (Figure 4B), with representative samples featuring a visibly expanded ECD and contracted pore (doi:10.5281/zenodo.4594193). Accordingly, closed as well as locally closed X-ray structures projected to the state-I cluster in all but the protonated wild-type system (Figure 4A). Two more states with contracted pores clustered at low–intermediate values of tIC1, varying somewhat with system conditions; these states were distinguished along tIC2 (high for state II, low for state III; Figure 4A). Representative conformations in state II were visibly similar to state I (doi:10.5281/zenodo.4594193); for the protonated wild-type system, the closed X-ray structure projected in State II, with locally closed structures along the state I–II border (Figure 4A). State III corresponded visibly to a partly expanded pore (doi:10.5281/zenodo.4594193), not represented by any known X-ray structures (Figure 4A). Open and lipid-modulated X-ray structures projected to a cluster at high tIC1 and low tIC2 (state IV, Figure 4A), substantially populated only upon protonation and/or I233T substitution (Figure 4B); as expected, representative samples featured the most expanded pore relative to other states (doi:10.5281/zenodo.4594193). A final cluster at high values of both tICs (state V, Figure 4A) was the least populated in all conditions (Figure 4B), again sampling intermediate values of pore expansion. Visual inspection of all four landscapes (Figure 4A) indicated that transitions from closed- (states I/II) to open-like (state IV) could proceed via state III (Figure 4A).

**Figure 4:**
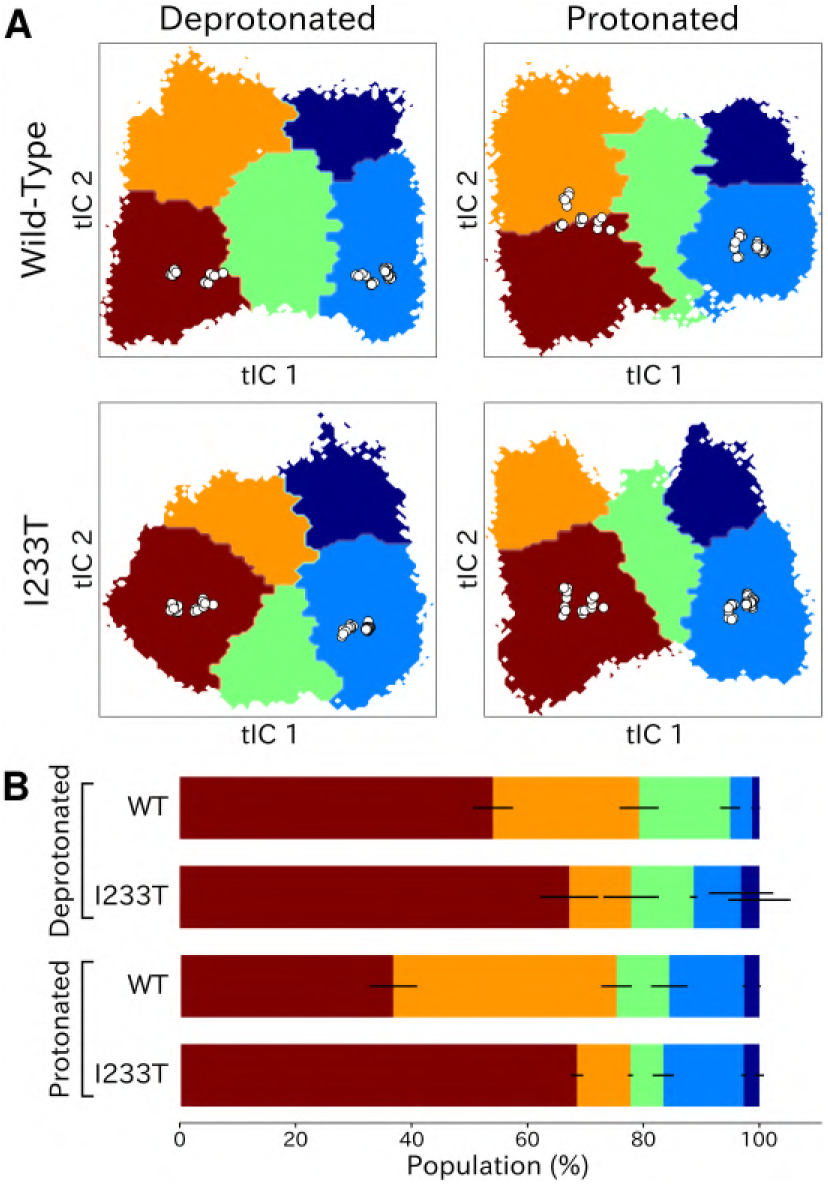
Higher-order clustering of the GLIC free energy landscapes. (A) Each free energy landscape can be further clustered into five-state models that - despite not being metastable - allow for more fine-grained structural analysis of the energy landscape. White dots represent experimental structures. (B) Populations for each of the five states. The following will be used to refer to the five regions: red - State I, orange - State II, green - State III, light blue - State IV, and dark blue - State V. Conformations sampled from these states can be accessed at doi:10.5281/zenodo.4594193.

To further validate this five-state space in context of past mechanistic models, we compared our clusters on the basis of conformational features implicated in channel gating. As previously described, a key distinction between closed- and open-like states was expansion of the upper-TMD pore, quantified here by the radial distance of the upper M2 helix from the pore center-of-mass (M2 spread). On this basis, states I/II and IV/V were respectively contracted and expanded, with state III sampling intermediate values (Figure 5A). The hydrophobic gate (9’, Figure S3A) also showed evidence of open-like state expansion. In addition, the pore radius at this position was generally expanded in mutant versus wild-type conditions (Figure S3A), as expected upon substitution of threonine for isoleucine. All states were somewhat constricted at 9’ relative to the open X-ray structure (Figure S3A), though our previous measurements confirmed sampling of hydrated conformations (≥ 30 water molecules in the channel pore, Figure 3D) in open-like states in both conditions. Interestingly, a proposed secondary gate at the intracellular end of the pore (−2’ radius, Figure S3B) implicated in desensitization [34] exhibited a bimodal distribution in radii, with state I largely retaining an expanded ≥5-Å radius similar to X-ray structures, while other states partially sampled a contracted radius - 2 Å.

**Figure 5:**
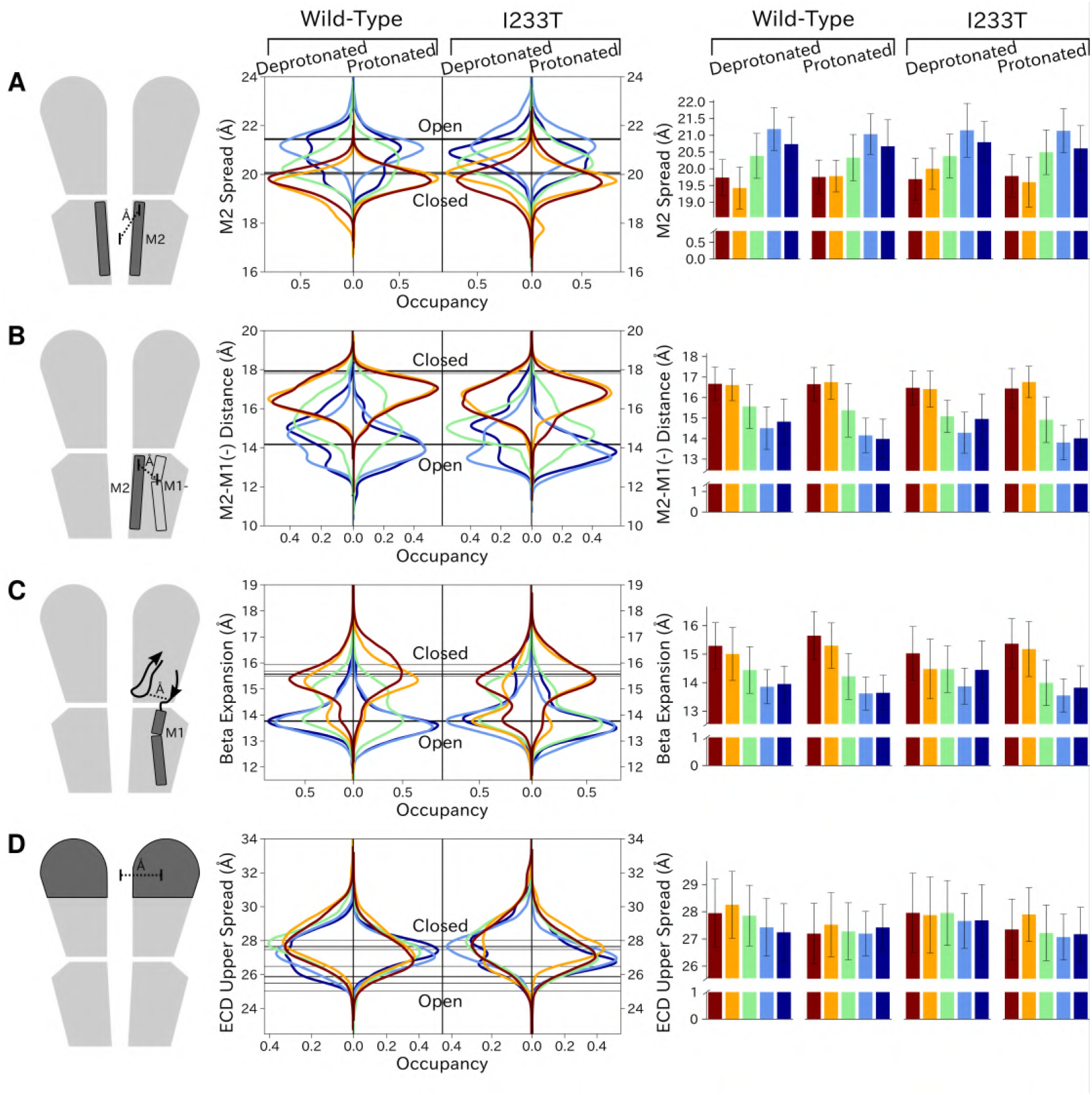
Probability distributions of a few variables proposed to be important in GLIC gating. The left-most cartoons illustrate the definition of each variable, while data is presented as probability distributions with means and standard deviations plotted as bars. Colors represent the five states in 4, and black lines represent the experimental structures 4HFI [27] and 4NPQ [26]. The spread of the pore-lining M2 helices (A) and the distance between the pore-lining M2 helix and the M1 helix of the neighboring subunit (B) are correctly captured by the 5-state model open and closed states, with intermediate states taking intermediate values. The beta expansion (C) yields distributions with expectation values of closed and open states aligning well with the experimental structures, while the intermediate state produces a bimodal distribution. Interestingly, the probability distributions of the closed-like states of the I233T mutation at deprotonated conditions show increased biomodality as well. The upper spread of the extracellular domain (ECD) (D) does not result in a clear separation of the five states, but a smaller pH-dependent shift can be observed.

In parallel with expansion of the upper pore, gating transitions have been associated with contraction at TMD subunit interfaces, quantified here by the distance between proximal regions of the principal upper-M2 and complementary upper-M1 helices (M2^+^-M1^−^ distance). Subunit interfaces in states I/II and IV/V were expanded and contracted respectively, with state III again sampling intermediate values (Figure 5B). Transitions at this interface have also been linked to relief of a helix kink, proximal to a conserved proline residue in M1. Indeed, this helix kink was more acute in states I/II, sampling even smaller angles than closed X-ray structures; conversely, the kink was largely relieved in state IV, with even larger angles than open structures (Figure S3C). State III again sampled intermediate values. Interestingly, kink angles for state V overlapped states I/II in deprotonated conditions, but shifted toward state IV in protonated conditions.

Gating transitions in the TMD are coupled to allosteric rearrangements in the ECD, particularly so-called *β*-expansion involving the first and last extracellular *β*-strands in each subunit. Proximal to the TMD interface, the cleft between these ECD strands is relatively expanded in closed X-ray structures but contracted in open structures, strengthening a salt bridge between *β*1-D32 and *β* 10-R192 [20]. Interestingly, values of *β*-expansion exhibited a bimodal distribution, with distinct peaks centered around closed and open structures (Figure 5C). In our MSMs, state-I samples were more expanded, while states IV/V were more contracted; distributions in states II/III featured distinct peaks at both closed- and open-like values.

More global gating motions in the ECD are thought to include blooming and twisting, with channel activation involving contraction and untwisting relative to the TMD. Surprisingly, no consistent state-dependent trends were noted in ECD spread (Figure 5D) or twist (Figure S3D). However, comparing e.g. the predominating state I in each condition revealed an overall pH dependence, with the ECD generally more contracted and twisted in protonated than deprotonated conditions (Figure 5D, Figure S3D). Interestingly, the ECD rarely contracted to the extent of open X-ray structures, nor twisted to the extent of closed X-ray structures in any condition, suggesting that crystal contacts may favor uncommon conformations in this domain.

### Symmetry Analysis Reveal Protonation- and State-Dependent Differences

Whereas each GLIC molecule is composed of five identical subunits, and exhibits five-fold symmetry in the context of a crystal, it remains unclear how symmetry is retained or broken in the course of channel gating. Previous simulations suggested a role for conformational asymmetry, particularly in the TMD, in facilitating closing transitions [4, 35]. To estimate subunit symmetry, we quantified pairwise RMSDs between homologous atoms in neighboring and opposing subunits, and plotted the resulting symmetry value for each simulation frame to its corresponding position in tIC space. In this representation, regions of higher pairwise RMSD correspond to lower symmetry. To enable identification of domain-specific changes, analyses were also performed independently for the TMD and ECD in each condition (Figure 6, top bars). In both protonated datasets, pairwise RMSDs of the open state were significantly lower compared to deprotonated conditions, suggesting that protonation plays a role in the symmetrization of this state. These high symmetry levels could be deduced from both the ECD and TMD (Figure 6B,D, top bars). Upon channel closure, the overall symmetry decreases, which is primarily driven by the symmetry loss seen in the ECD. The TMD on the other hand seems to recover some symmetry in the closed state. At deprotonated conditions, pairwise RMSDs were generally higher across the board, and neither ECD or TMD were able to reach the low RMSD levels of the open state at protonated conditions. The ECD symmetry still seemed to decrease upon channel closure, but the differences were diminished compared to protonated conditions (Figure 6A,C, top bars).

**Figure 6:**
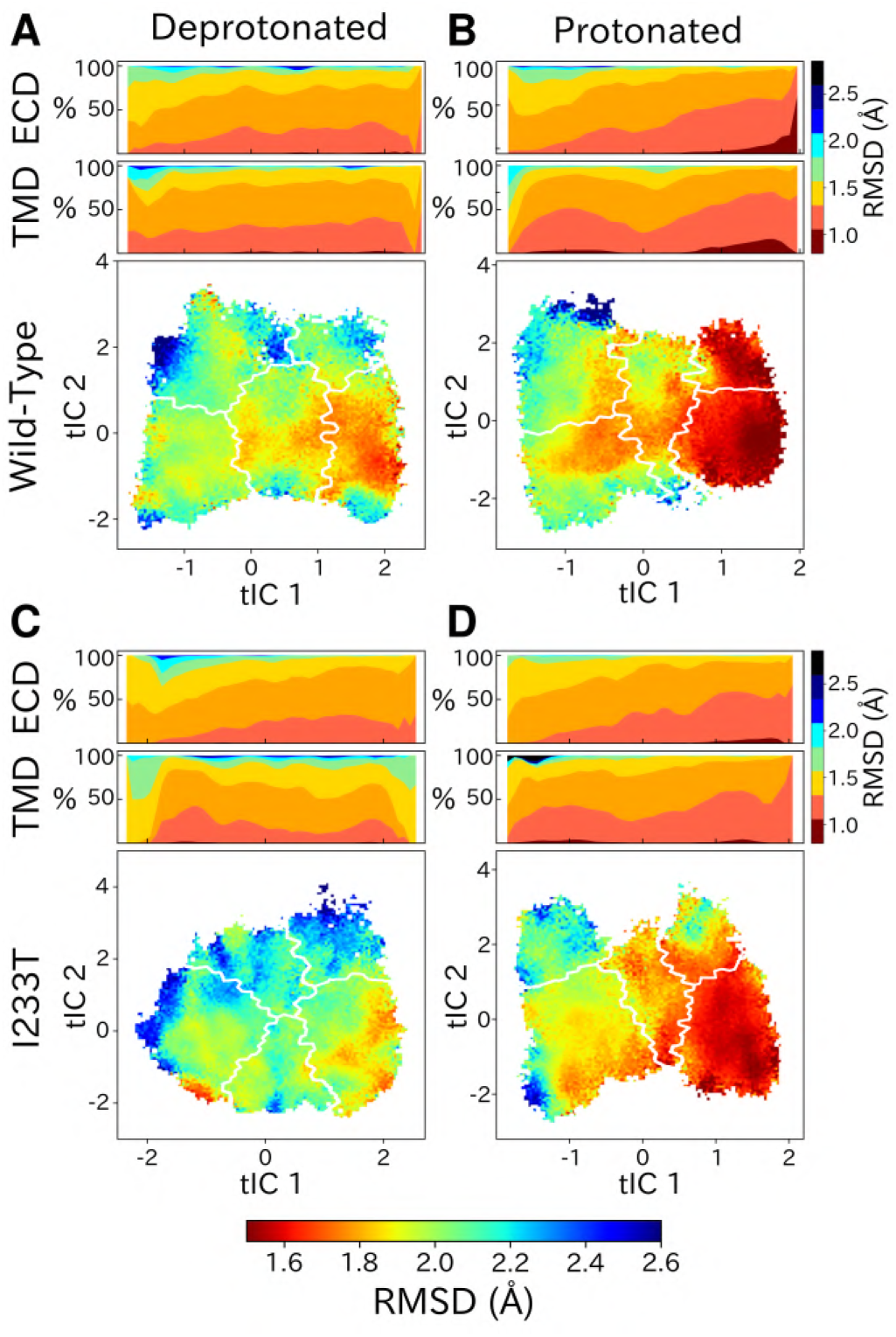
State- and protonation-dependent differences in ion channel symmetry. Heatmaps show pairwise RMSDs between all subunits of the channel, measuring the conformational symmetry of the pentamer. The two top bars show pairwise RMSDs of the transmembrane (TMD) and extracellular (ECD) domains separately, represented as stacked histograms along tIC 1. At deprotonated conditions wild-type (A) and the I233T mutation (C) show decreased levels of symmetry in both TMD, ECD and overall. At protonated conditions wild-type (B) and the I233T mutation (D) display high levels of symmetry in the open state coming from both ECD and TMD. Notably, the symmetry of the ECD is decreasing during channel closure and total closed state symmetry is coming mainly from the TMD.

## Discussion

We have constructed Markov state models of a pentameric ligand-gated ion channel that enabled quantitative modeling of protonation and mutation effects, identification of intermediate states and characterization of the effect of symmetry in channel gating. Our free energy landscapes showed deepening of the open state well upon protonation and destabilization of the closed state upon mutation of the hydrophobic gate (Figure 7A), but at any point in time only a minority fraction of channels will actually be open, in agreement with electrophysiology recordings. In addition to capturing features of the gating mechanism already proposed for pLGICs, our MSMs allowed for further exploration of features that correlate with gating. Here, we focused on the effect of conformational symmetry between subunits and found that the open state displayed higher levels of symmetry, which was particularly enhanced after protonation (Figure 7B).

**Figure 7:**
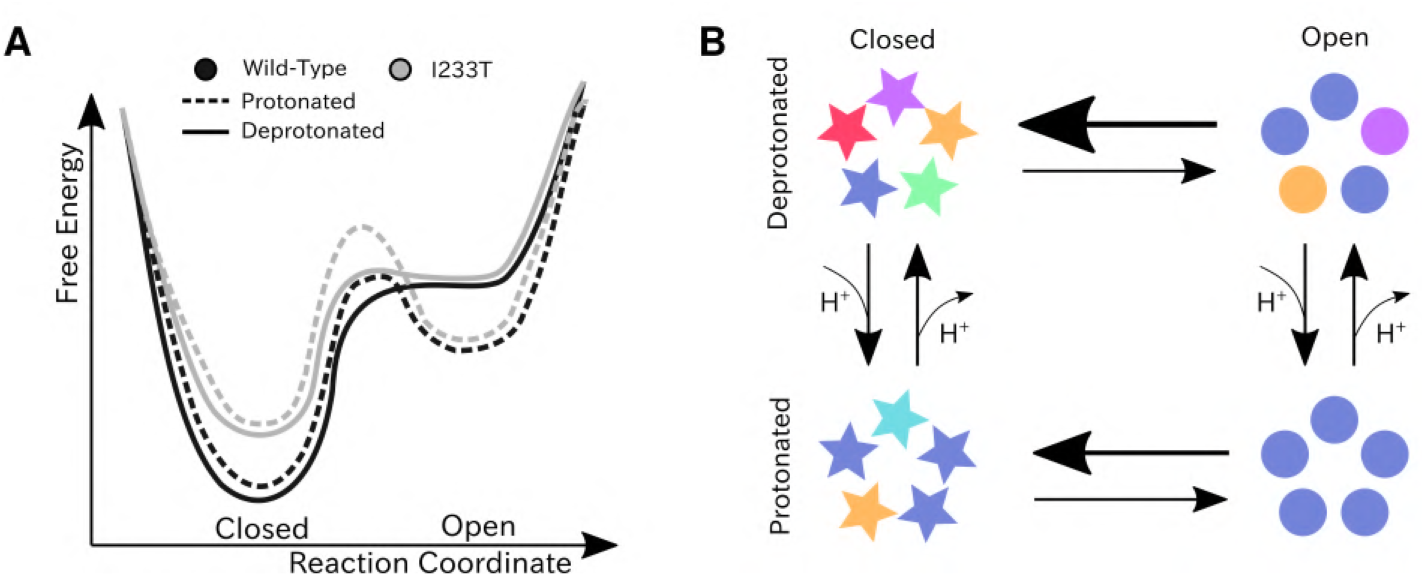
Proposed models for the free energy landscape and symmetrization in GLIC gating. (A) A sketch of the free energy landscapes for the wild-type and I233T mutant at protonated and deprotonated conditions. An open-state well is formed when the channel is protonated and the I233T mutation destabilizes the closed state, but only a small fraction of channels will be open at any point. (B) Conformational symmetries of GLIC are affected by the protonation state. Upon protonation the open state displays a high level of symmetry, which is reduced to an intermediate level in the closed state. When deprotonated, the open state achieves an intermediate level of symmetry, which is reduces to low levels of symmetry in the closed state. This suggests that protonation is important for symmetrization of the open state.

Thermodynamic properties calculated from the present models were largely consistent with functional recordings, showing a shift towards relative stabilization of open versus closed states upon protonation of acidic residues or polar substitution at the 9’ hydrophobic gate (Figure 7A). Free energies of the gating transition have previously been estimated using string method optimization of a few collective variables assumed to be important in the gating transition [20]. The variable associated with the largest barrier height in the work of Lev et al. is pore hydration, which is an integral part of the gating transition, thus suitable for comparison to the maximal energy barrier height from the MSMs. At low pH the string method hydration yielded a barrier of 1.5 kcal/mol, compared to 1.0-1.5 kcal/mol for the protonated MSM. At neutral pH the barrier from string method hydration gave 2.5 kcal/mol, while the MSM resulted in 1.5-2.0 kcal/mol height of the energy barrier. This indicated that the wild-type MSMs found transition pathways with slightly lower free energy barriers.

Two-state clustering suggested a value of P_open_ well below 100% for GLIC, even at activating conditions. Although P_open_ has not been determined for GLIC, recent cryo-EM structures of closed-state GLIC solved at activating conditions point towards a significant closed population even at activating conditions [26, 33]. Other channels in the pLGIC family have also been shown to attain a range of maximal open probabilities; 10-40% for GABA_A_Rs [36, 37], 20-80% for 5-HT_3_s [38, 39], 0.2-3% for nAChRs [40, 41], and 90-100% for GlyRs [42, 43], when saturated with their respective natural agonist. Given the spread in open probability between pLGIC subtypes it seems reasonable that a more distant bacterial homolog like GLIC could have a unique energy landscape. Additionally, since the open probabilities for GLIC are less than 100%, protonation could function like partial rather than full agonism.

Five-state clustering enabled more detailed investigation of the different regions of the energy landscapes, including intermediate and alternative open-like and closed-like conformations. In all cases, open and modulated crystal structures projected in state IV, while closed and locally-closed states projected into state I, or on the border between state I and state II for the protonated wild-type dataset. Locally closed structures projected in the same free energy basin as closed-state 4NPQ but closer to the activation free energy barrier, indicating that the locally closed state could serve as a pre-activating state in the gating pathway. Prevost *et al.* solved crystal structures of multiple locally closed states trapped by various mutations, and out of these only one was capable of opening in electrophysiology experiments. Additionally, a range of conformations in the M2-M3 region were sampled in the cryo-EM structures [7], indicating that the locally closed conformations might not represent a separate metastable state in wild-type GLIC.

Structural studies of the open, closed and locally closed crystal structures have revealed several conformational changes associated with GLIC gating [44]. The first steps are thought to occur in the ECD through blooming and twisting motions of the entire domain. Surprisingly, our simulations did not capture statedependent but rather pH-dependent differences of these motions, suggesting that the ECD blooming and twisting motions happen quickly in response to a change in pH. Conversely, we did observe transitions in the loops connecting the inner and outer sides of the ECD *β* sandwich, which have been shown to be important for pLGIC channel function. In particular, crystal structures show breakage of the D32-R192 salt bridge in closed-state GLIC and mutational studies of D32 reveal loss of function [45]. In addition to capturing state-dependent differences in agreement with experimental data, our analysis also showed bimodality of the probability distributions at neutral pH for the closed states (Figure 5). This effect was further enhanced upon mutation of the 9’ gate, indicating that there could be an allosteric pathway between the center of the transmembrane pore and the D32-R192 salt bridge. This is also supported by previous computational models based solely on the apparent open structure [46].

Gating motions in the TMD are particularly characterized by the tilting of the pore-lining M2 helices, leading to constriction around the 9’ hydrophobic gate followed by pore dehydration. This process has previously been observed in early simulations of the TMD by Zhu and Hummer [1–3]. Our models successfully captured how these M2 helix motions closely correlated with changes in 9’ radius and pore hydration levels (Figures S12, S13, 3D). Even though we capture state-dependent differences in the 9’ pore radius, and individual simulations sample pores that are as wide as the open-state structures, the most probable conformations of our open state clusters do not exhibit pores as expanded as in the majority of open crystal structures (Figure S3). These structure have typically been co-crystallized with a hydrophobic plug of detergent molecules between the 9’ region and the top of M2, associated with a more expanded, funnel-shaped form, which is hardly sampled in simulations after plug removal. Other variables that have been associated with GLIC channel gating, including kinking of the M1 helix [44] and interactions in the TMD-TMD subunit interface [47], were also captured by our MSMs (Figure S3C, Figure 5B). Notably, our observations were largely consistent with [20], validating the use of these features as collective variables. Interestingly, the protonated alternative open-like state V displayed particular constriction at the −2’ gate incompatible with ion conduction (Figure S3). This is a typical feature of desensitized states in the pLGIC family [34], although a desensitized state for GLIC has not yet been resolved. Energetically, open and desensitized states would be expected to be separated by an energy barrier, which is not the case in our models, but it is possible state V represents conformations relevant for pre-desensitization.

Analysis of the conformational symmetry of GLIC revealed a particularly symmetric open state at protonated conditions (Figure 6). At deprotonated conditions the symmetry levels were lower overall, indicating that protonation is important for channel open-state symmetrization. In all cases, significantly less symmetric ECDs could be observed in the closed state compared to the open state, although the difference was further enhanced at protonated conditions due to the more symmetric open state. The vast majority of GLIC structures cover the open state and display high levels of ECD symmetry, while the fewer structures covering the closed state are relatively poorly resolved, possibly indicating heterogeneity particularly in the ECD [33, 44]. These results suggests that ECD symmetry could serve as an entropic driving force, where perturbation to the symmetric structure of the open state ECD facilitates channel closure. At activating conditions the ECD becomes protonated which facilitates high-level symmetrization characteristic for the open state. Furthermore, at protonated conditions the TMD displayed increased levels of symmetry both in the open state and closed state with less symmetric conformations in between. At deprotonated conditions, the symmetry level of the TMD transition was largely consistent with that of the protonated conditions, but the TMDs of the closed-like conformations were less symmetric. So far consensus has not been reached regarding whether homomeric channels transition symmetrically or asymmetrically, but evidence points towards asymmetric transitions being common. Recent cryo-EM structures of the 5-HT_3A_ receptor in lipid nanodiscs revealed an asymmetric closed state, whereas a symmetrized open state was stabilized by serotonin molecules bound at all five ECD sites [48]. Additionally, long (15-20 μs) simulations of 5-HT_3A_ have shown evidence of asymmetric closure of the TMD pore upon channel pre-activation [49]. Mowrey et *al.* also proposed that asymmetric propofol binding could create unbalanced forces such that symmetry breaking would facilitate channel conformational transitions [35]. In our simulations, ligands (protons) were bound symmetrically across all subunits which further indicates that asymmetric transitions could be important regardless of symmetric or asymmetric ligand-binding.

In summary, our Markov states models are able to predict shifts in free energies upon change in activating stimulus (pH) and upon mutation. The models predict relatively low values of maximal open probability, with relative differences in agreement with electrophysiology recordings. Our simulations also captured state-dependent differences in previously proposed mechanistic variables as well as state- and pH-dependent differences in channel conformational symmetry. These models allow for further exploration of conformational changes that correlate with channel gating to improve understanding of the gating mechanisms of pLGICs, which is important both for fundamental understanding of channel gating and the development of state-selective drugs.

## Materials and Methods

### Pathway Construction Using eBDIMS

Seeds along the GLIC closed-open gating pathway were obtained from 50 models generated along forward and reverse eBDIMS simulations [28, 29]. The channel was represented as an elastic network model using the C*α* representation of apparent closed (PDB ID 4NPQ) [26] and open (PDB ID 4HFI) [27] structures, and driving transitions in both directions by progressively minimizing the difference in internal distances between the current and the target states. Langevin dynamics with implicit solvent and harmonic forces was used to model the dynamics, with a 12 Å intrasubunit and 8 Å intersubunit cutoff, respectively. The force constant of the elastic network was 10 kcal/(mol·Å), as previously used in [28].

### Model Reconstruction

Atomistic detail of the seeds was reconstructed using Modeller version 9.22 [50] in two steps. First, side chain atoms from the template X-ray structure (PDB ID 4HFI) were added to each model, followed by a cycle of refinement with all C*α* atoms restrained. Restraints on C*α* atoms were then substituted with restraints on backbone hydrogen bonds, taken from helix and sheet annotations in the template PDB file, for another cycle of refinement.

### Molecular Dynamics Simulations

The reconstructed seed models were embedded in a palmitoyloleoylphosphatidylcholine (POPC) bilayer, solvated by water and 0.1 M NaCl. Activating conditions were modeled by protonation of a subset of acidic residues (E26, E35, E67, E75, E82, D86, D88, E177, E243; H277 doubly protonated) to approximate the probable pattern at pH 4.6, as previously described [30]. All systems were energy minimized with steepest descent for 10,000 steps. NPT equilibration was carried out in four steps, restraining all heavy atoms in the first cycle, then only backbone atoms, then C*α* atoms, and finally the M2 helices, for a total of 76 ns. Equilibration and production runs were performed using the Amber99SB-ILDN force field with Berger lipids, together with the TIP3P water model. Temperature coupling was achieved with velocity rescaling [51] and pressure coupling with the Parrinello-Rahman barostat [52]. The simulations were prepared and run with GROMACS versions 2018.4 and 2019.3 [53] for 1.2 *μs* each.

### Markov State Models

The GLIC ion channel was described with a set of 1585 features, including interatomic distances within and between subunits. Since GLIC is a symmetric homopentamer, we introduced a feature transformation where all distances originating from one subunit were scored as an independent trajectory; thus each trajectory contained information as to how one subunit moved in relation to all others. Dimensionality reduction was achieved using tIC analysis, previously shown to form a good basis for discretization of conformational space into Markov states, as the tIC eigenvetors linearly approximate the MSM eigenvectors [54, 55]. Typically, MSMs are constructed with the assumption of exhaustive sampling of the equilibrium distribution, and inclusion of as much kinetic information as possible. Although this is a feasible approach for peptide-sized systems, we find it practically unfeasible for large-scale motions in ion channels since the MSM tends to optimize for slow but undersampled processes. Accordingly, we discretized the state space into a few, but meaningful, tIC dimensions by omitting faster dimensions where the data was represented as a single Gaussian. Hyperparameter optimization is commonly solved by maximizing the variational approach for Markov porcesses (VAMP) [56] score with cross-validation to avoid overfitting; however, in our case we found that VAMP favored explorative behavior rather than convergence of a few interesting processes in the absence of of exhaustive equilibrium sampling. Instead, we relied on a simpler elbow approach, where open probabilities were calculated from a PCCA+ [57] two-macrostate model for various hyperparameter combinations. Based on the resulting plots (Figure S6), we selected 300 clusters for kMeans clustering, noting the results were consistent for other hyperparameter combinations. The tIC lag times yielded consistent results within a 5-25 ns range for all datasets (Figure S5), so we selected a lag time of 20 ns, with kinetic mapping [58]. The MSM lag time was determined to 20 ns from the implied timescales (Figure S4). One simulation from the protonated wild-type dataset was rejected from analysis due to obstruction of the tIC landscape with a slower but undersampled process, leaving 58.8 *μs* total sampling; MSMs the other three conditions (deprotonated wild-type, deprotonated and protonated I233T) contained 60 μs sampling each. Markov state modeling was done with PyEMMA version 2.5.7 [59].

### Electrophysiology

Two-electrode voltage-clamp electrophysiology was performed as previously described [12]. Briefly, GLIC cDNA subcloned in vector pMT3 was modified using commercially synthesized primers (Invitrogen, Stockholm, Sweden) and the GeneArt site-directed mutagenesis system (Thermo Fischer, Waltham, MA). Plasmids were amplified using a HiSpeed Plasmid Purfication Midi kit (Qiagen, Hilden, Germany), and verified by cycle sequencing (Eurofins Genomics GmbH, Ebersberg, Germany). Nuclei of isolated stage V–VI *Xenopus laevis* oocytes (EcoCyte BioScience, Dortmund, DE) were injected with 0.5–3.0 ng cDNA and stored in incubation medium (88 mM NaCl, 10 mM HEPES, 2.4 mM NaHCO3, 1 mM KCl, 0.91 mM CaCl2, 0.82 mM MgSO4, 0.33 mM Ca(NO3)2, 2 mM sodium pyruvate, 0.5 mM theophylline, 0.1 mM gentamicin, 17 mM streptomycin, 10,000 u/L penicillin, pH 8.5) for 2–7 days. Glass electrodes were pulled and filled with 3 M KCl to reach an initial resistance of 0.1-0.5 MΩ. Expressing oocytes were clamped at −70 mV using an OC-725C voltage clamp (Warner Instruments, Hamden, CT, USA), and perfused with running buffer (123 mM NaCl, 10 mM HEPES, 2 mM KCl, 2 mM MgSO4, 2 mM CaCl2, pH 8.5) at a flow rate of 0.35 mL/min. Activation buffers contained 10 mM MOPS or citrate in place of HEPES, adjusted in 0.5-pH unit increments, and were exchanged for running buffer via a VC^3^-8 valve controlled pressurized perfusion system (ALA Scientific Instruments, Farmingdale, NY, USA). Currents were digitized at a sampling rate of 5 kHz with an Axon CNS 1440A Digidata system using pCLAMP 10 (Molecular Devices, Sunnyvale, CA, USA). Each pH response was measured as the peak current after 1 minute exposure to activation buffer, normalized to the same oocyte’s response to the lowest pH tested. Each reported value represents the mean from 6-8 oocytes, ±standard error of the mean. Proton concentration-dependence curves were fit by nonlinear regression with bottom and top restraints using Prism 8.0 (GraphPad Software, La Jolla, CA).

## Acknowledgements

This work was supported by grants from the Knut and Alice Wallenberg Foundation, the Swedish Research Council (2017-04641, 2018-06479, 2019-02433), the Swedish e-Science Research Centre, and the BioExcel Center of Excellence (EU 823830). Computational resources were provided by the Swedish National Infrastructure for Computing. The authors are grateful to Joseph Jordan, Brooke Husic, and Lucie Delemotte for helpful feedback and discussions.

## Data Availability

Additional data including simulation parameters and sampled conformations from the MSMs can be accessed at doi:10.5281/zenodo.4594193. Due to size limitations full trajectories are available upon request.

## Supporting Information

**Figure S1:**
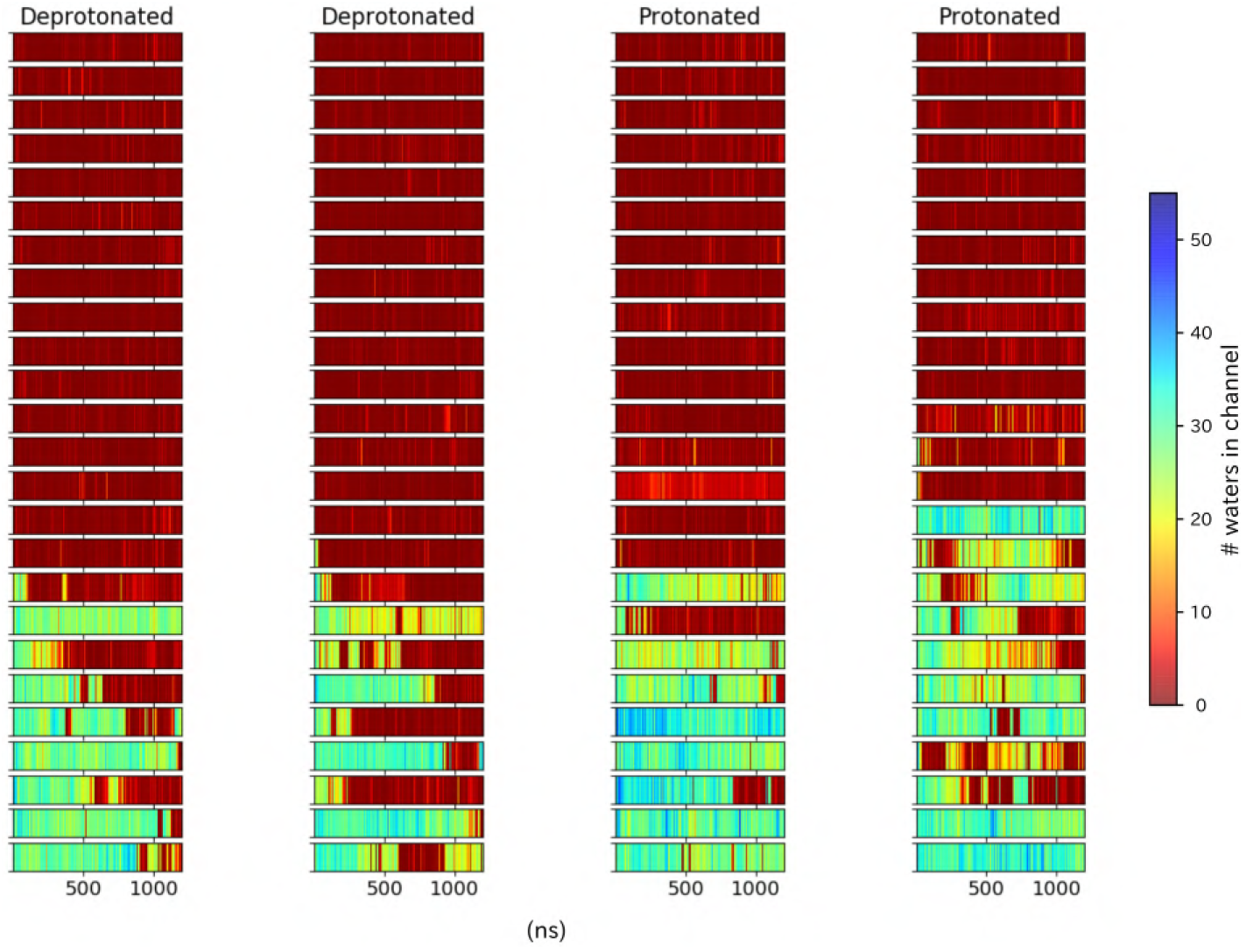
Pore hydration in wild-type GLIC simulations. Hydration of all 100 wild-type simulations, where each row represents one simulation with frames on the x-axis. The simulations have been grouped based on location of the starting seed in the transition, with open-like seeds at the bottom and closed-like seeds at the top.

**Figure S2:**
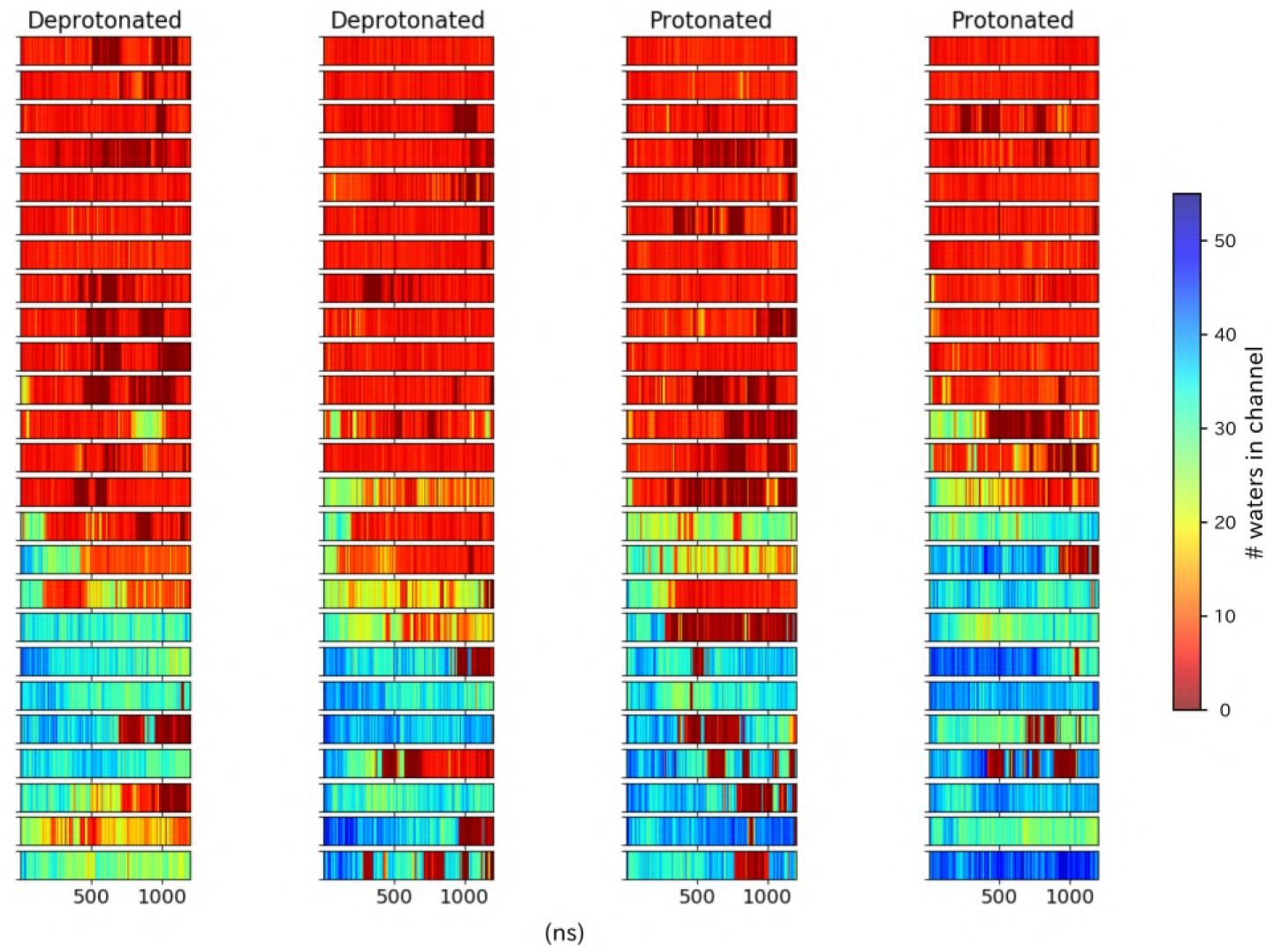
Pore hydration in I233T mutant GLIC simulations. Hydration of all 100 I233T simulations, where each row represents one simulation with frames on the x-axis. The simulations have been grouped based on location of the starting seed in the transition, with open-like seeds at the bottom and closed-like seeds at the top.

**Figure S3:**
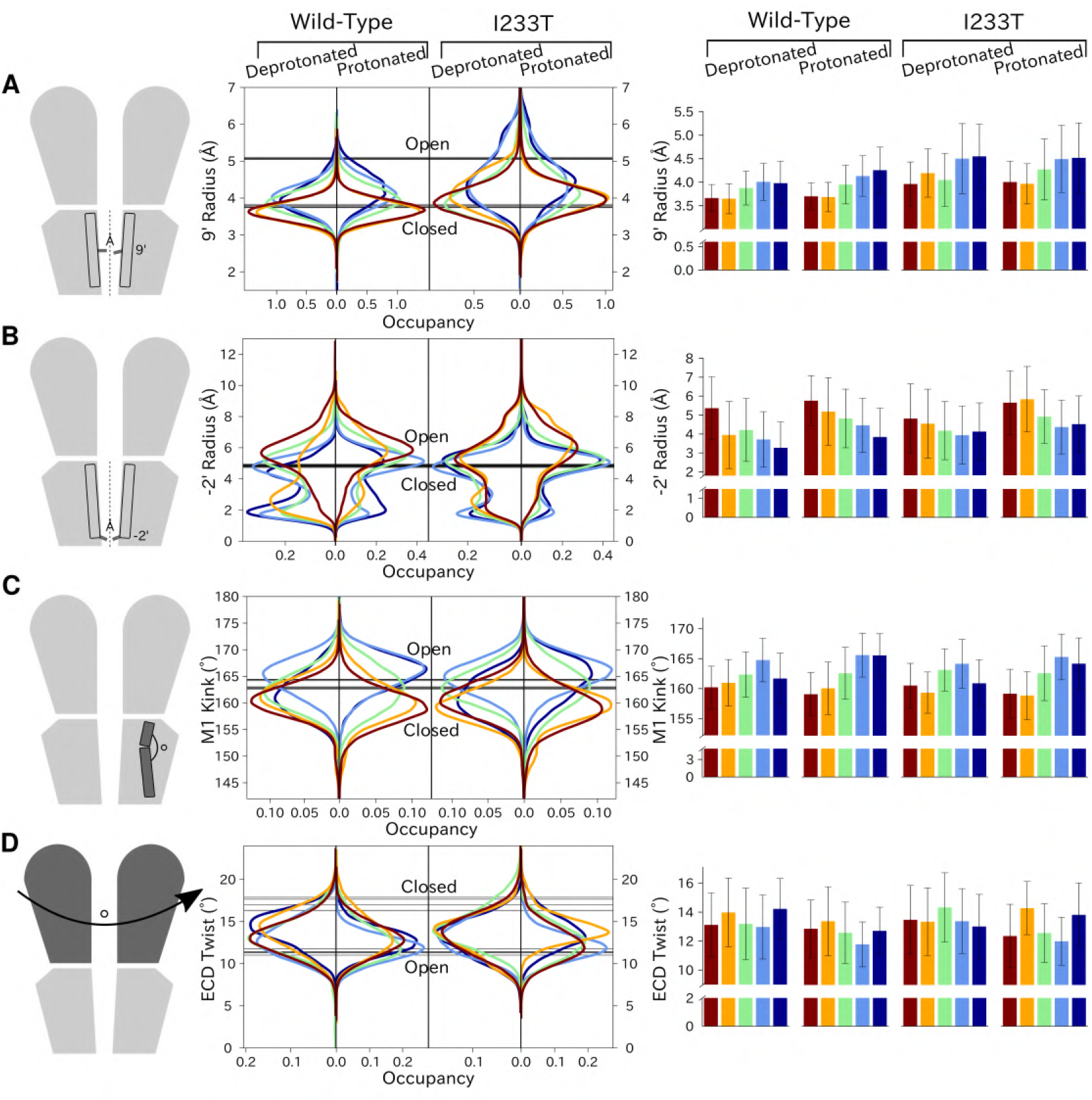
Probability distributions of a few variables proposed to be important in GLIC gating. The left-most cartoons illustrate the definition of each variable, while data is presented as four probability distributions with means and standard deviations plotted as bars. The experimental X-ray structures 4HFI and 4NPQ are shown as black lines together with the probability distributions. Colors represent the different states in 4. Kinking of the M1 helix is more pronounced when approaching the closed state (A). The twisting of the extracellular domain (B) does not result in a clear separation of the different states, but a small pH dependent shift can be observed. The radius from the pore center to the 9’ hydrophobic gate (C) is higher in the open states for all datasets. We note that the open X-ray structure, 4HFI, is more expanded compared to the simulations, probably due to the presence of a hydrophobic plug which was removed in the simulations. We also note that the I233T mutation results in a significantly more expanded pore. The radius between the pore and the −2’ gate (D) is lower in the open states for all datasets although the values for the closed and open experimental structures are the same.

**Figure S4:**
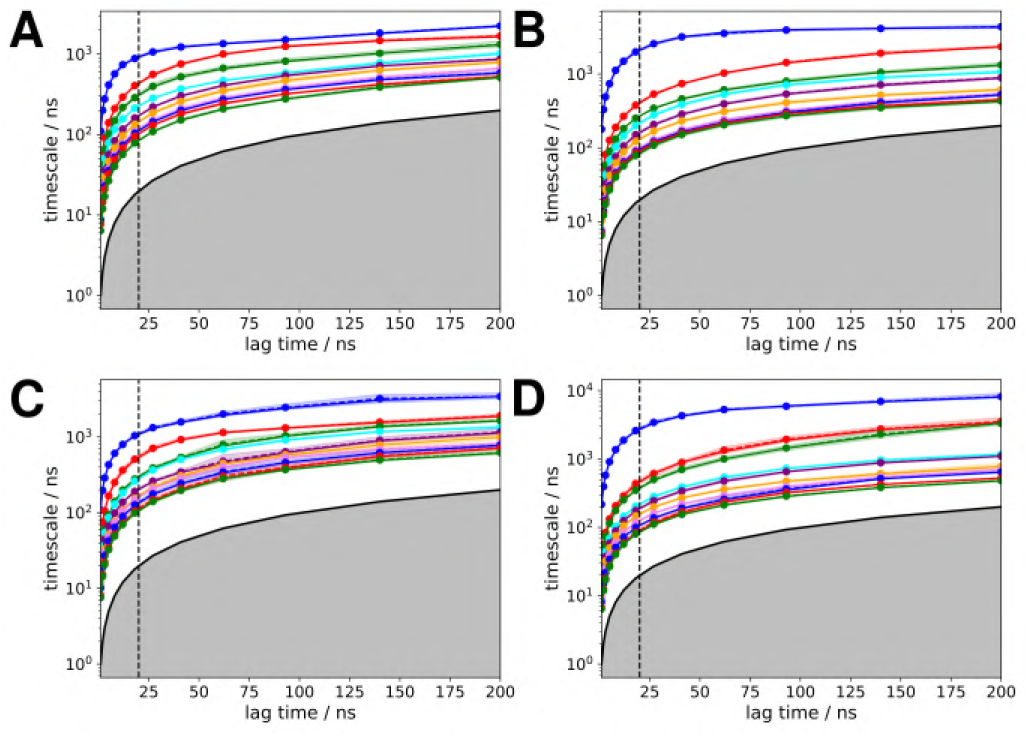
Implied timescales as a function of MSM lag time. The first 10 implied timescales representing the 10 slowest processes of the MSM, for (A) deprotonated wild-type, (B) protonated wild-type, (C) deprotonated I233T, (D) protonated I233T. The vertical dotted line represents the selected MSM lag time, chosen as the smallest lag time where the implied timescales have leveled out.

**Figure S5:**
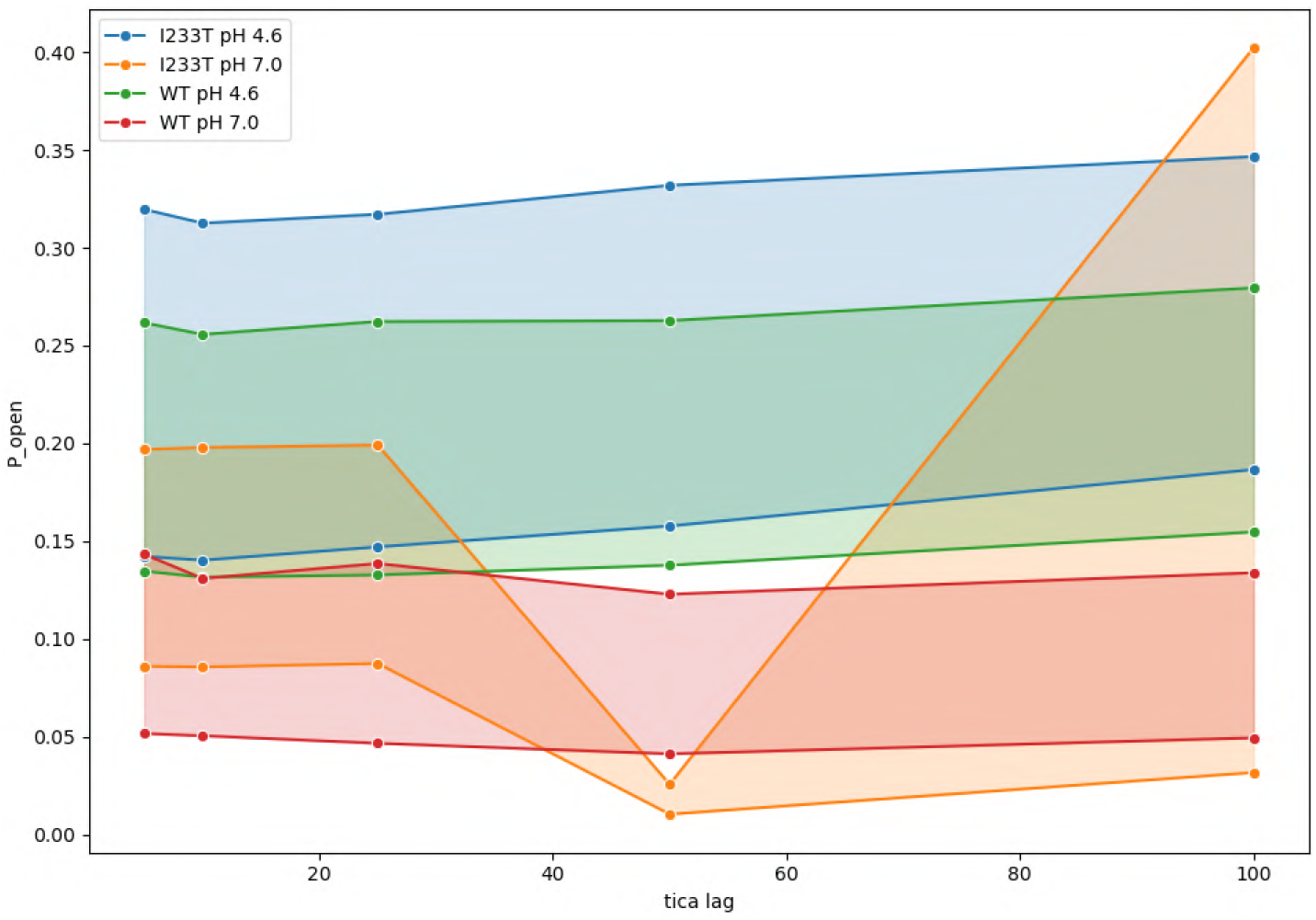
Open probabilities for all 2-state models as a function of tICA lag times. The width of each line represents other hyperparameter combinations, including number of microstate clusters and commute or kinetic mappings. The results are constant for all four datasets up to tICA lag time of at least 25 ns.

**Figure S6:**
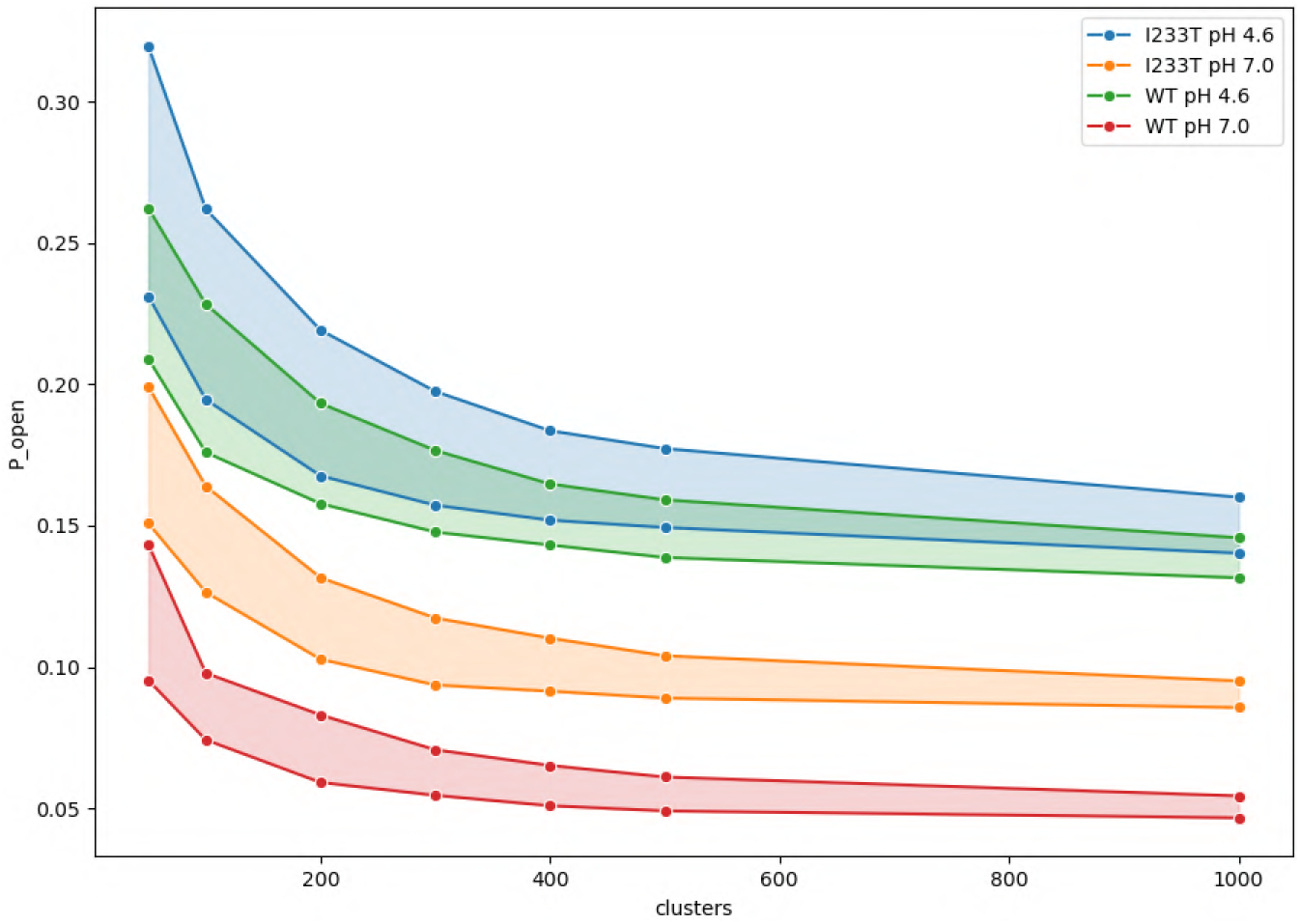
Open probabilities for all 2-state models as a function of the number of microstate cluster. The width of each line represents other hyperparameter combinations, including tICA lag times up to 25 ns together with either commute or kinetic mappings. 300 clusters were selected as the smallest number of clusters where the open probabilities had leveled out.

**Figure S7:**
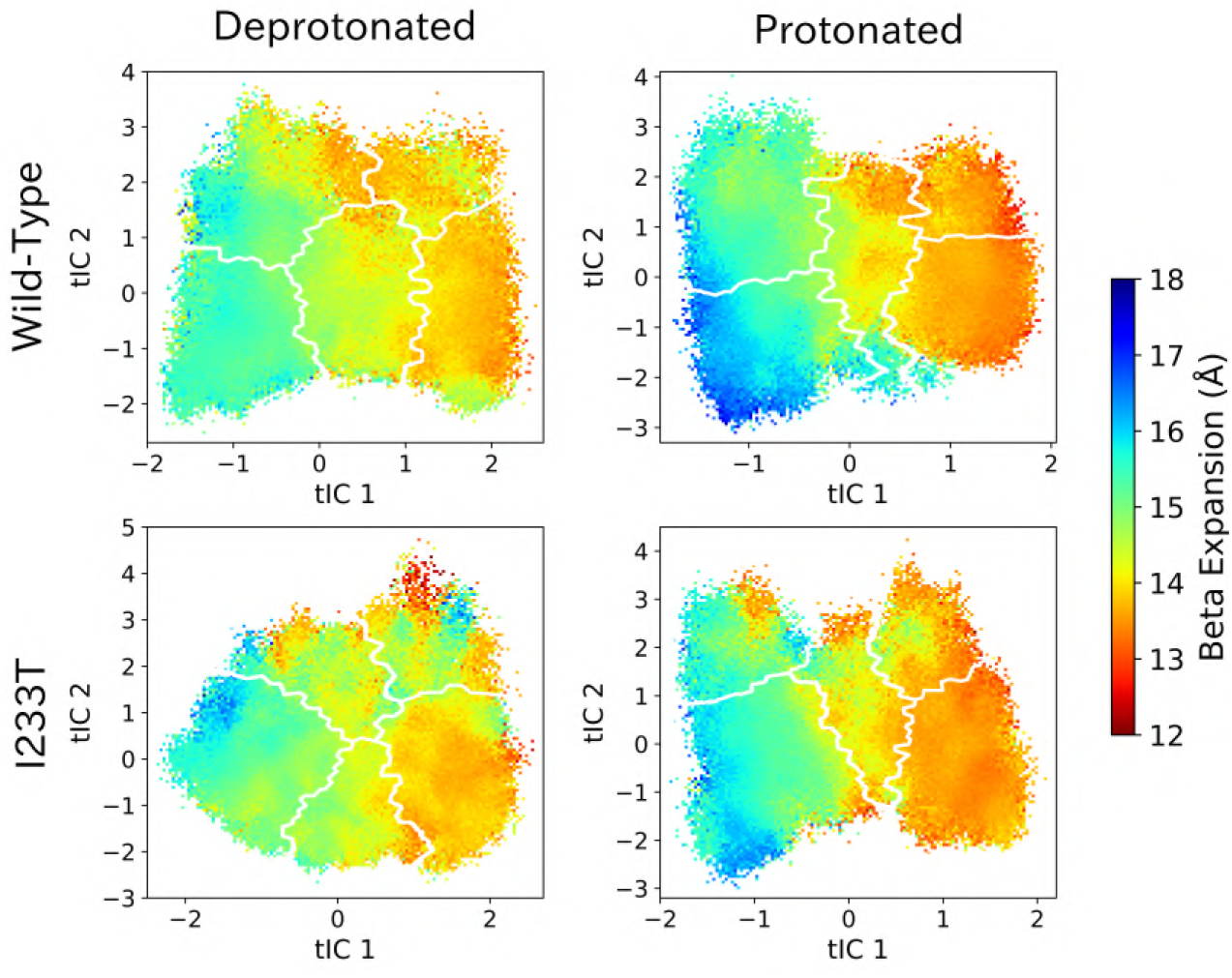
Beta expansion values projected onto the first two tICs. Line contours represent the five states in Figure 4.

**Figure S8:**
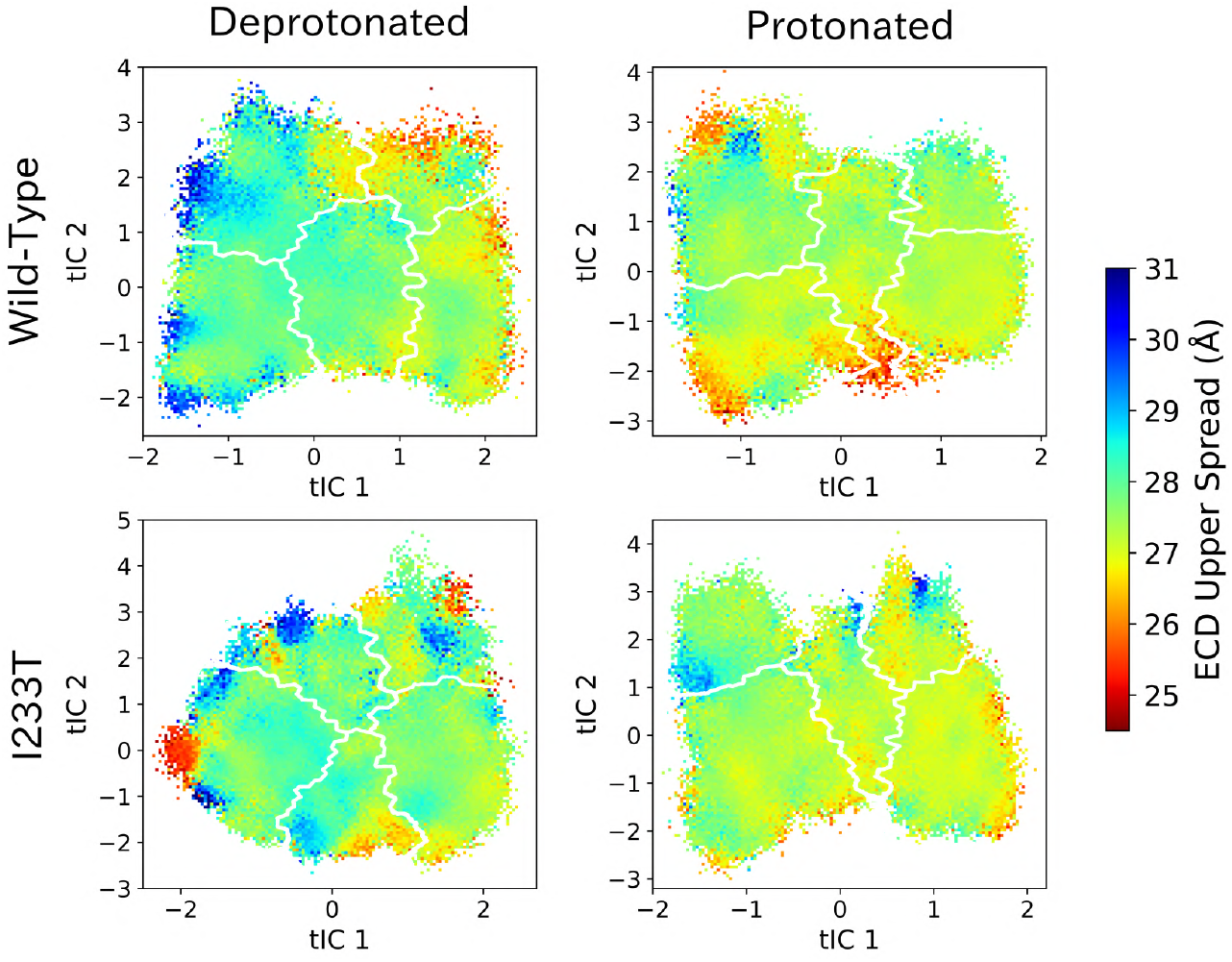
ECD upper spread values projected onto the first two tICs. Line contours represent the five states in Figure 4.

**Figure S9:**
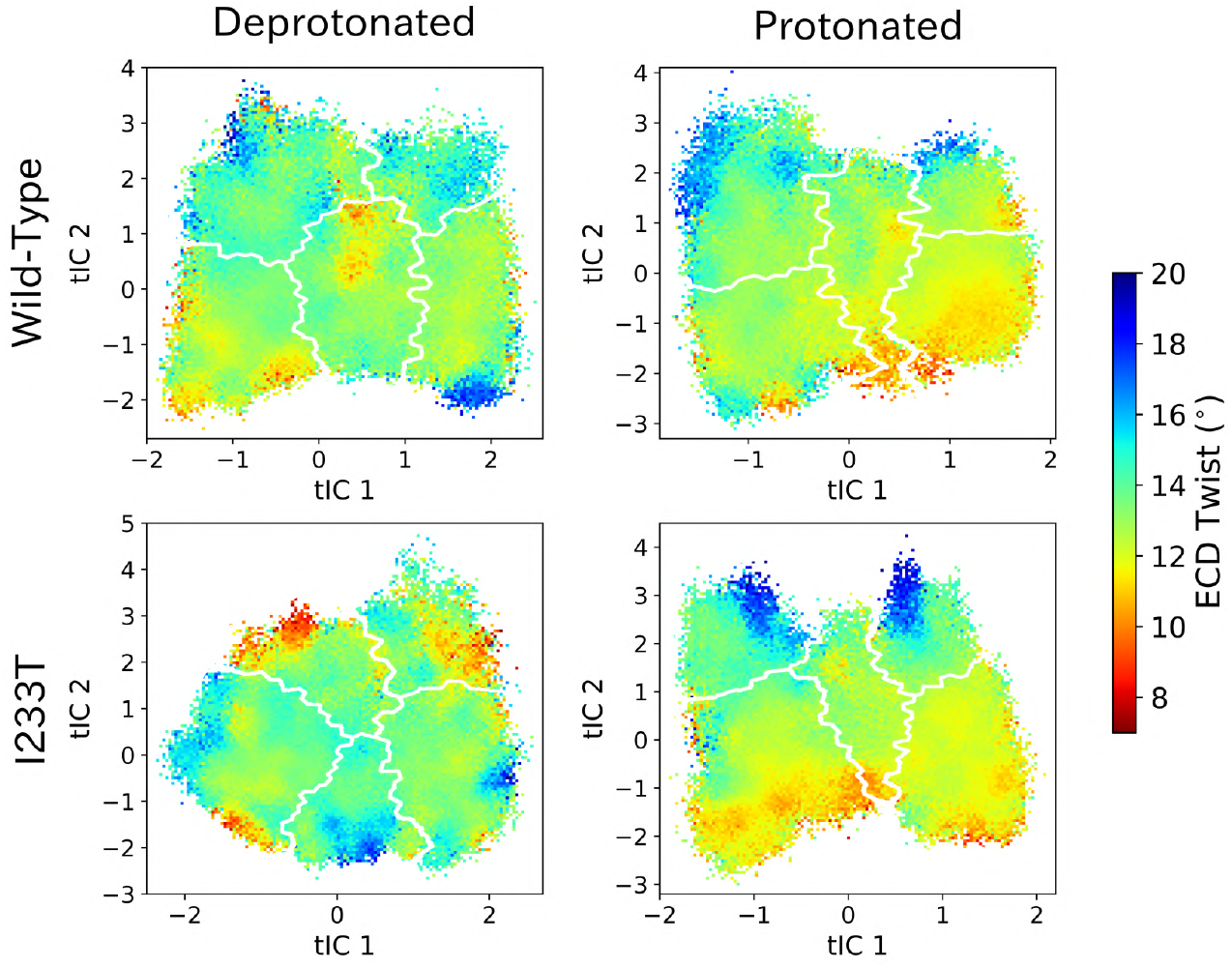
ECD twist values projected onto the first two tICs. Line contours represent the five states in Figure 4.

**Figure S10:**
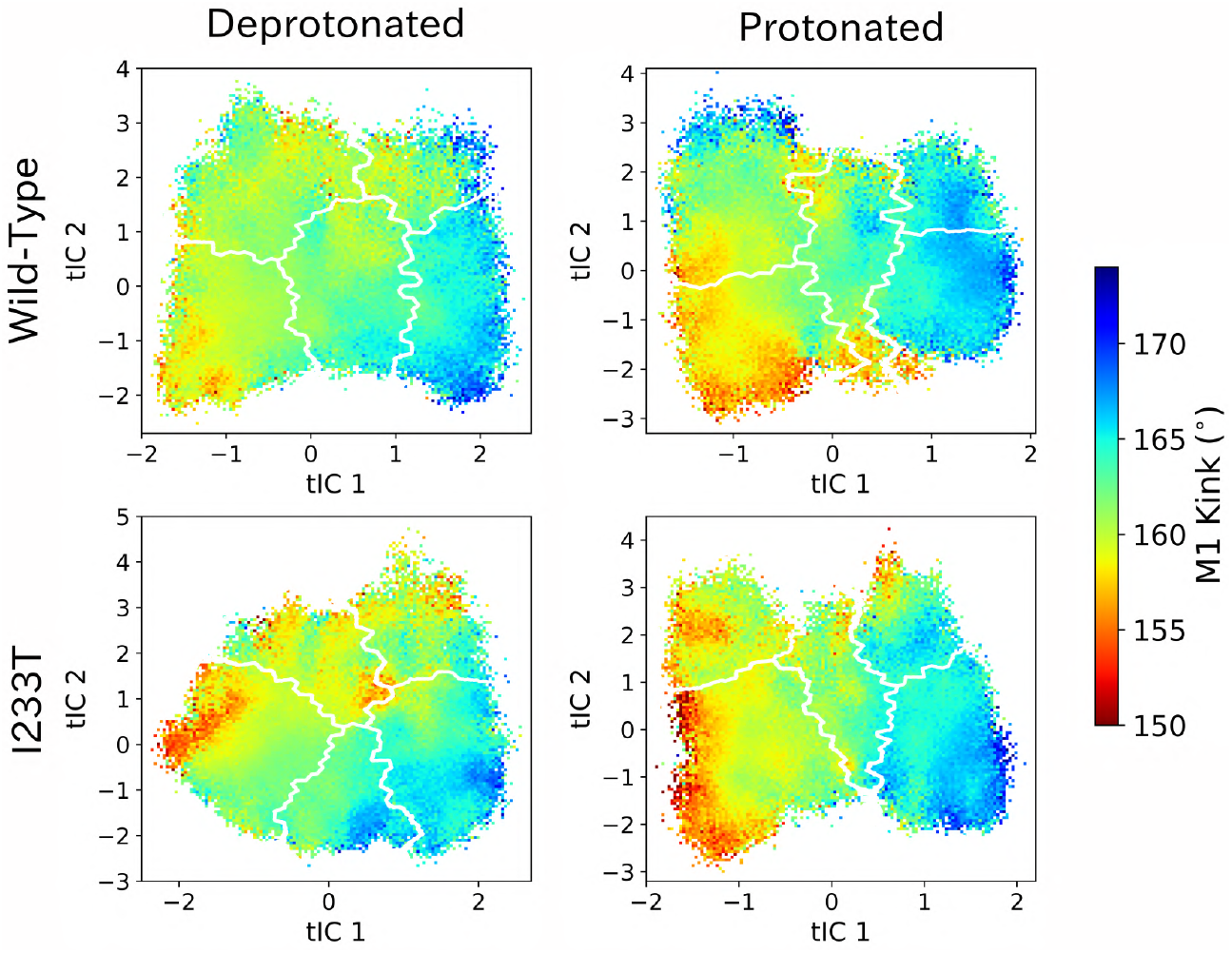
M1 kink values projected onto the first two tICs. Line contours represent the five states in Figure 4.

**Figure S11:**
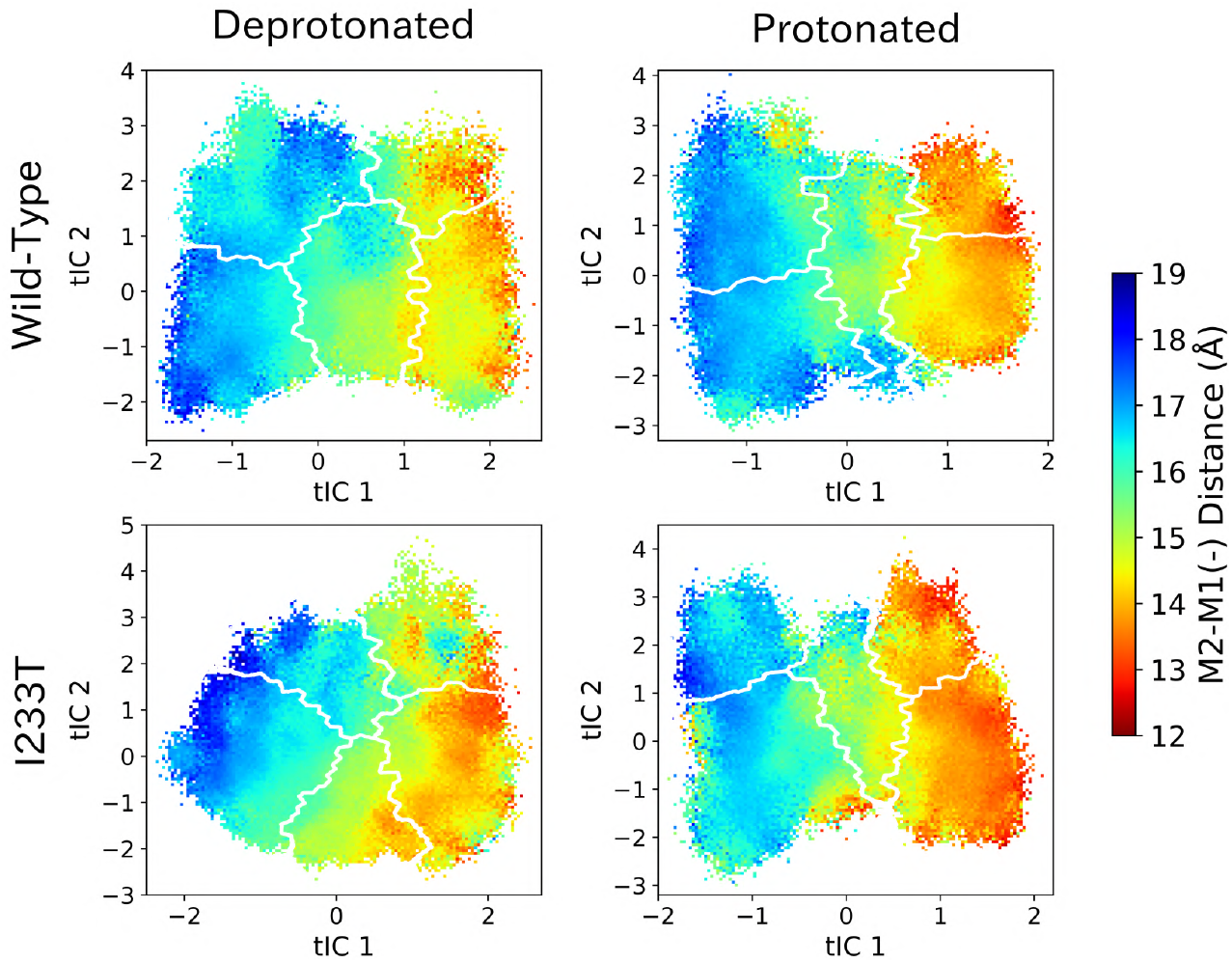
M2-M1(-) values projected onto the first two tICs. Line contours represent the five states in Figure 4.

**Figure S12:**
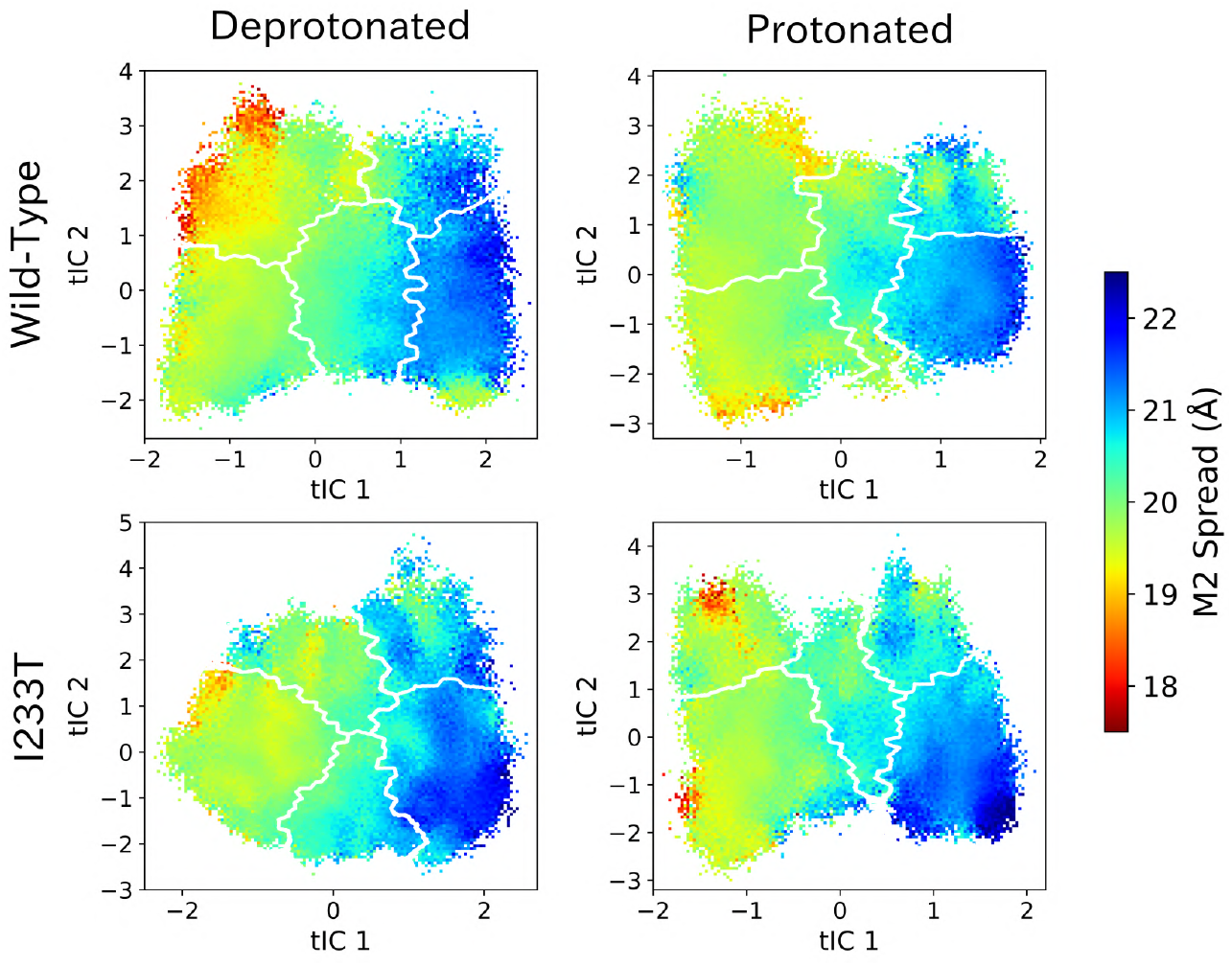
M2 spread values projected onto the first two tICs. Line contours represent the five states in Figure 4.

**Figure S13:**
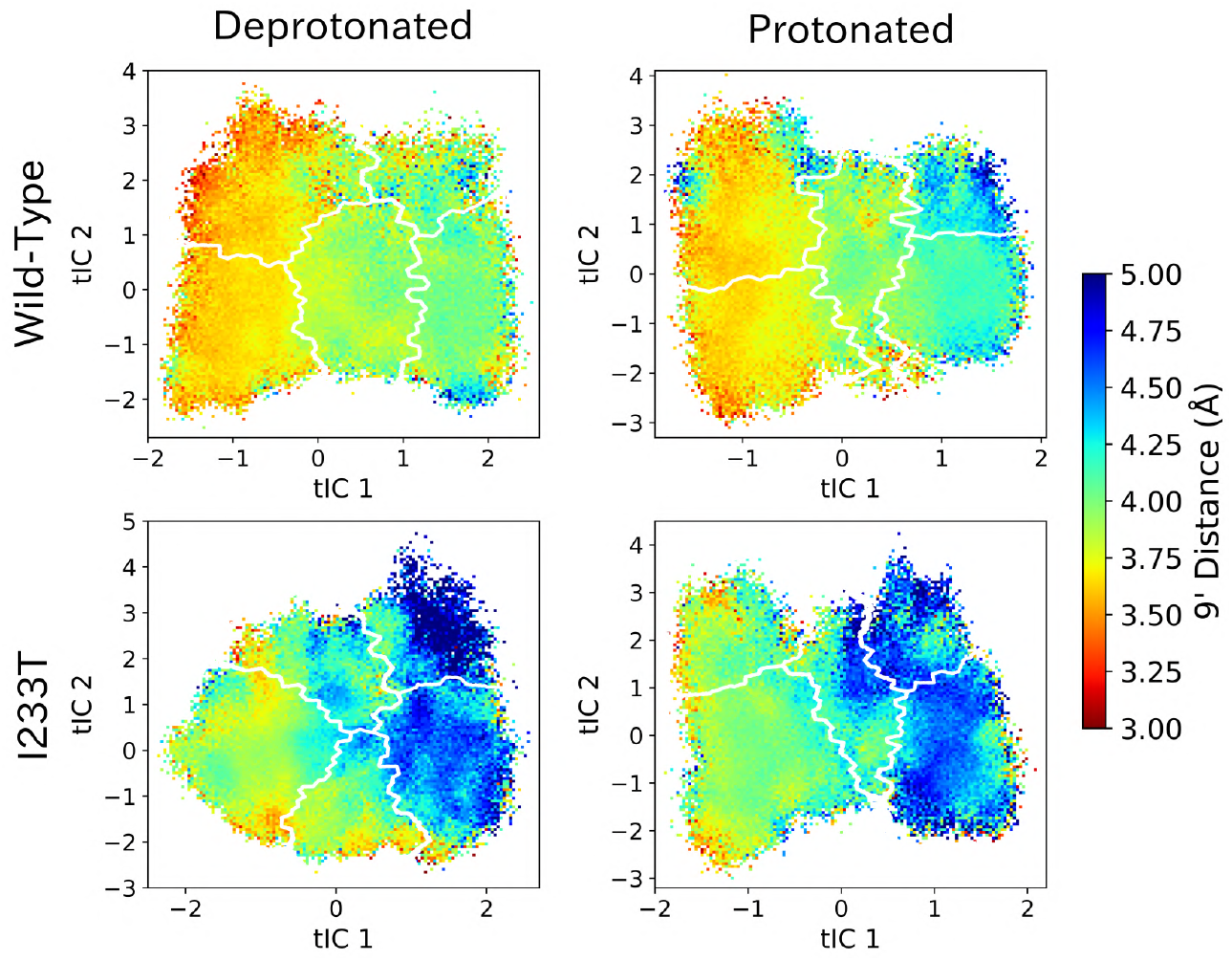
9’ distances projected onto the first two tICs. Line contours represent the five states in Figure 4.

**Figure S14:**
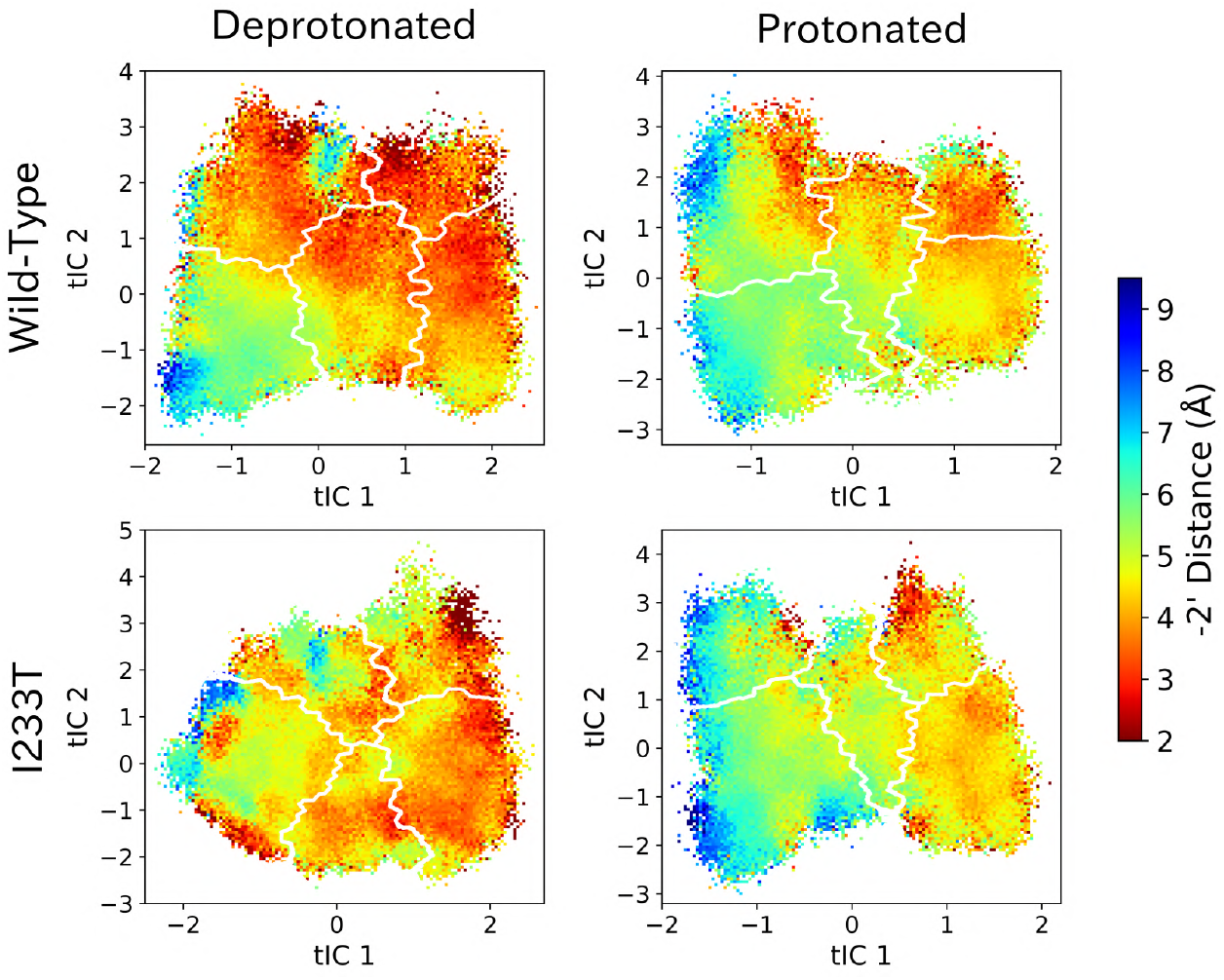
-2’ distances projected onto the first two tICs. Line contours represent the five states in Figure 4.

